# Bacterial peptide deformylase inhibitors induce prophages in competitors

**DOI:** 10.1101/2025.06.06.656871

**Authors:** Julie Chen, Katarina Pfeifer, Kerrin Steensen, Anne Marie Crooke, Michael Wolfram, Jacob Bobonis, Anna Lopatina, Nina Bartlau, Fatima A. Hussain, Emily P. Balskus, Martin F. Polz, Paul C. Blainey

## Abstract

While antibiotics mediate chemical warfare among microbes, their roles in the wild extend beyond direct growth inhibition(*1*). Some antibiotics have the potential to mediate interference competition by triggering a bacterial stress response that subsequently activates endogenous viruses integrated in bacterial genomes (prophages). Canonically, this activation is regulated by the SOS response upon DNA damage. Here we show that a metabolite produced by natural isolates of *Vibrio ordalii* circumvents the SOS response by directly triggering prophage induction in other *Vibrio* species, co-occurring in the same environment. While the metabolite was previously classified as a broad-spectrum antibiotic, we observe how it acts as a peptide deformylase inhibitor that specifically induces certain prophages, even when target bacterial cells carry multiple other prophages. Its biosynthetic gene cluster, or *ord* cluster, also encodes its own peptide deformylase (OrdE) which provides self-immunity to producer strains. Likewise, among natural *Vibrio* isolates that carry similar prophages, resistance against the *ord* metabolite was found in those that had acquired a divergent second peptide deformylase. Finally, we show that prophage induction by the *ord* cluster prevents slower-growing producer strains from being outcompeted by their otherwise fast-growing competitors if they carry an inducible prophage. Thus, we demonstrate how natural products play additional impactful roles in communities beyond antibiotic activity and that prophage induction serves as an interference competition strategy, sustaining community diversity.

---

Prophages are highly abundant mobile genetic elements (MGEs) found within bacterial genomes(*2*, *3*). When dormant, these viruses passively proliferate with the host genome, even endowing the host with superinfection exclusion/immunity proteins, or additional metabolic functions(*4–6*). Still, prophages present risks to their hosts as timebombs that can destroy the cell when they are induced to replicate. Such induction may be triggered remotely by natural compounds produced by other microbes, making prophages a powerful target for interference competition (in contrast to the indirect inhibition by consumption of shared resources). Prophage induction has been linked to the SOS stress response system where the SOS system is activated in response to DNA damage. The host RecA protein then also stimulates the autoproteolysis of the phage cI repressor, resulting in host machinery being co-opted by the phage through prophage induction(*7*, *8*). In fact, the primary approach to prophage discovery has been to treat cells with high concentrations of potent DNA damaging agents like Mitomycin C (MMC) or ciprofloxacin that are known to activate the SOS system. Cases of prophage induction by environmental competitors have also been linked to the SOS response via the production of hydrogen peroxide or colibactin(*9–11*). However, it is now clear that DNA damage does not encompass the diversity of induction cues in nature, and that induction screens relying on DNA damaging agents do not capture all inducible prophages(*12*). Recent examples support the existence of SOS-independent mechanisms of prophage induction by exogenous chemicals including direct induction by autoinducers(*13*, *14*), the signaling molecule in quorum sensing, and pyocyanin(*15*), a redox-active phenazine pigment, further demonstrating the potential of non-canonical inducing agents and pathways. Based on this evidence, we reasoned that a broad screen for interference competition, designed to detect phage-mediated cell lysis between natural isolates, may reveal additional mechanisms of prophage induction.

To discover cases of prophage induction triggered by other bacteria, we developed a high-throughput combinatorial microfluidic screening pipeline aimed at first detecting cell lysis when bacteria are co-cultured and then verifying which of such cases are associated with prophage induction. Our screening focused on a collection of 110 bacterial strains of the genus *Vibrio* all isolated from the same coastal marine environment in Massachusetts. Using subsets of our collection, previous studies have demonstrated antagonism mediated by secondary metabolites, interactions between MGEs/phages and their hosts, as well as prophage diversity(*16–19*). We further interrogated putative cases of prophage induction in screened co-cultures using a combination of molecular genetics and comparative genomics, leading to the characterization of an antagonistic interaction mediated by SOS-independent, selective prophage induction exhibited by a pair of isolates had displayed the highest lysis and viral induction levels. We found that induction was controlled by a metabolite that acts as a peptide deformylase inhibitor. We also showed that the host can be rescued from lysis by prophage induction upon the expression of a horizontally acquired peptide deformylase with low similarity to the genomic peptide deformylase.

## Results

### Antagonism screen for prophage induction

To maximise throughput of our pairwise antagonism screen, we implemented the kChip - a droplet microfluidics screening platform that enables the massively parallel assembly and measurement of combinations of isolates(*20*, *21*). The kChip allowed us to perform 120,000 pairwise microscale co-cultures in parallel per device, including controls. Thus, we were able to assay 5,995 unique isolate pairs covered by 24,530 conditions (four relative abundance ratios per co-culture and the respective monocultures, see Materials and Methods), powered to a median of 14 replicate co-cultures (fig. S1A). To detect cellular lysis during co-culture, we implemented a fluorescence-based time course assay in the kChip using the non-toxic dye SYTOX Green which intercalates DNA but cannot permeate intact cell envelopes (fig. S1B-D). To confidently score lysis due to interactions, we calculated lysis scores by comparing the signal from co-culture to that of a baseline comprising the combined respective monocultures, then scaling the resulting “lysis scores” from -1 to 1 where positive scores denote co-cultures with signals greater than the respective monocultures (Materials and Methods). The utility of this lysis assay was confirmed by comparing it to the traditional Burkholder plate-based assay using isolates known to induce lysis during co-culture(*15*) (fig. S1E-F). We recapitulated the co-cultures that displayed high lysis in the kChip (Materials and Methods) using a similar fluorescence assay in microtitre plates. Co-cultures exhibiting lysis at both assay scales were then evaluated for prophage induction by sequencing pooled samples and searching for an increase in coverage in genomic regions that exhibited sequence characteristics consistent with prophages (Materials and Methods). These cases were verified by quantifying phage production with phage-specific qPCR assays (Materials and Methods).

Our screen found that 2.9% (172 of 5,995) of all unique pairwise co-cultures lead to lysis by antagonism with the distribution skewed towards six strains that were involved in lytic interactions with ≥25% of the collection (fig. S2, data file S1). From these 172 co-cultures, our sequencing-based screen identified ten putatively induced phages from eight isolates; these phages were also all part of the 107 independently and bioinformatically predicted prophages within our 110-isolate collection (Materials and Methods, data file S2). To delineate spontaneous induction from direct induction by a specific isolate, we individually co-cultured each of these eight hosts with each isolate that led to a positive lysis score, then measured induction by comparing viral loads using phage-specific qPCR between co-cultures and their corresponding host monocultures. Altogether, three phages reached high titres relative to monoculture, prompting us to hypothesize that induction occurred due to antagonism for: phage 41 in *Vibrio cyclitrophicus* 1F-97, phage 5.64 in *Vibrio tasmaniensis* FF-266, and phage 6.141 in *Vibrio ordalii* FF-93, while the remainder displayed high spontaneous induction (data file S1). All three phages are found in polylysogenic strains and have small genomes (33.6 kbp - 37.5 kbp), where phage 41 additionally appears to be a circular, episomal prophage with an estimated four copies per host genome. The three phages also varied in the number of bacterial isolates that triggered their induction in co-culture, the genetic distances between these isolates, as well as the degree of induction (fig. S3). Our study focuses on the induction of phage 41 which displayed the highest lysis and relative viral load in our screen when its host, *V. cyclitrophicus* 1F-97, was specifically co-cultured with *V. ordalii* isolates 12-B09 or FS-144, whereas the other *V. ordalii* strains did not induce lysis (fig. 1A-B, fig. S4A). Using 12-B09 as a model hereon, we confirmed the directionality of the interaction with the Burkholder assay (fig. S4B). With the strong response of phage 41 in *V. cyclitrophicus* 1F-97 confirmed, we sought to determine the mechanism of induction.

**Fig. 1.**
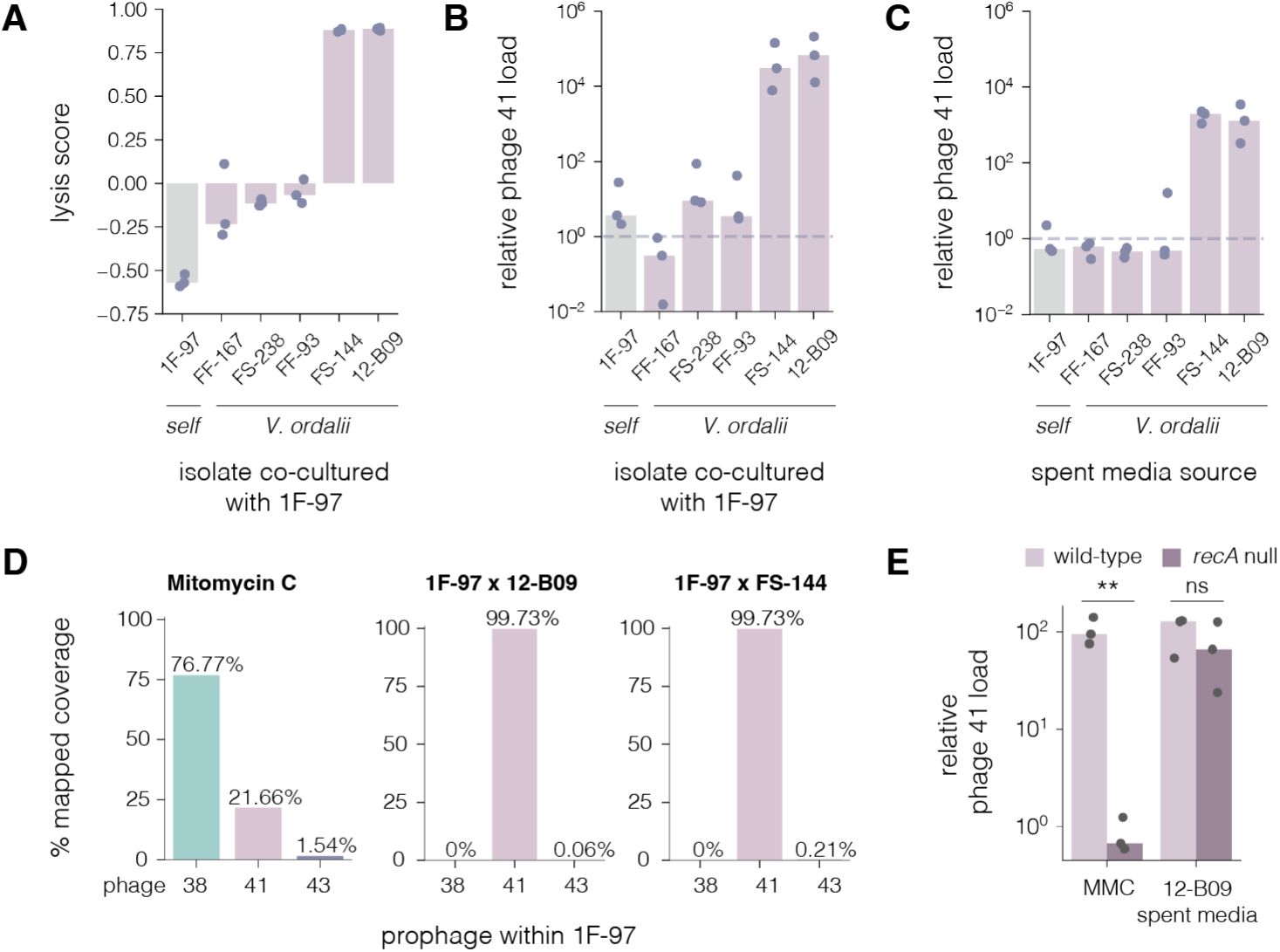
*V. ordalii* isolates specifically induce phage 41 in 1F-97 in an SOS-independent manner. (A) Lysis scores for co-cultures comprising *V. cyclitrophicus* 1F-97 and all *V. ordalii* isolates in the collection or itself, all at starting inoculum of OD_600_ 0.01 (0.01 x 0.01). Self (1F-97) cultures differ from monoculture in that a second 1F-97 culture is combined in place of a *V. ordalii* culture. Positive scores indicate lysis by co-culture. (B,C) Estimated induction (relative viral load) of phage 41 of strain 1F-97 when (B) co-cultured with or (C) treated with the spent media from each of the *V. ordalii* isolates or itself for 24-h. (D) Percentage of coverage from concordantly mapped reads for the prophages carried by 1F-97 (phages 38, 41, and 43) under the following conditions: 1F-97 treated with 0.05 µg/mL Mitomycin C during logarithmic phase (left) or co-cultures (0.01 x 0.01) comprising 1F-97 with either *V. ordalii* isolates 12-B09 (middle) or FS-144 (right). The final mapped read counts: 14.3 M (MMC), 1.8 M (12-B09 co-culture), 9.2 M (FS-144 co-culture). The resulting median coverage across the entire phage regions in order of phages 38, 41, and 43: 80,825/25,834/1,892 (MMC); 0/15,663/10 (12-B09 co-culture); 0/78,962/174 (FS-144 co-culture). (E) Estimated induction of phage 41 of either 1F-97 strains, wild-type or the *recA* null mutant, after treatment by 0.05 µg/mL Mitomycin C (MMC) induced at OD_600_=0.2-0.35 or at OD_600_=0.45-0.6 in equal parts with 12-B09 spent media. Asterisks (**) indicate p-value <0.01 calculated using a two-sided t-test, ns stands for “not significant” (p-value <0.05). (A) Lysis score bars represent the mean of three biological replicates from 22-h time courses in 384W microtitre plates with individual replicates shown (mean of four technical replicates). (B,C,E) Bars represent the median of three biological replicates where each replicate is the mean of three technical qPCR replicates (also shown as individual points). (B,C) Dashed line represents no difference (1X) from the untreated 1F-97 culture.

### SOS-independent induction of phage 41

First, we evaluated whether the antagonism by *V. ordalii* 12-B09 and FS-144 is contact-dependent or -independent. To that end, we asked whether induction is effected by a secreted metabolite by treating 1F-97 with nutrient-supplemented spent media from both inducing and non-inducing *V. ordalii* isolates. Similarly to the co-culture experiments, induction was only observed with 12-B09 and FS-144 spent media (fig. 1C). This suggests that induction is neither contact-dependent, nor due to compounds generally present in spent media, but rather is due to a specifically secreted product likely encoded by a biosynthetic gene cluster (BGC) unique to 12-B09 and FS-144.

*V. cyclitrophicus* 1F-97 is a polylysogenic strain containing two previously described prophages: an SOS-responsive phage (phage 38) and a putatively dual SOS and quorum-sensing responsive phage (phage 43)(*22*), and also a third prophage which is our focus here (phage 41). Intriguingly, we observed that *V. ordalii* isolates 12-B09 and FS-144 triggered the nearly exclusive induction of phage 41 in an SOS-independent manner (fig. 1D). To understand the differences in induction between these three prophages, we compared the relative phage population levels across various inducer types (fig. 1D). While all three prophages are induced when exposed to the DNA damaging agent Mitomycin C (MMC), phage 38 dominates the other two by up to an order of magnitude. In contrast, only phage 41 is induced when 1F-97 is co-cultured with 12-B09 or FS-144, while the other two prophages were close to background induction levels. To confirm whether the induction of phage 41 is independent of the canonical stress response, we constructed a *recA* null mutant in 1F-97 that abolishes the SOS response. Despite the loss of the RecA-dependent SOS system, 12-B09 spent media could induce phage 41, while induction was lost upon MMC treatment (fig. 1E). Furthermore, the lack of observable phage 41 induction in pure 1F-97 culture suggests that spontaneous induction is low (raw reads deposited on Zenodo, see Data and materials availability). Altogether, the specific and SOS-independent induction of phage 41 in 1F-97 can be triggered by select *V. ordalii* strains or their spent media, raising further questions about a potentially novel class of prophage inducers.

### A hybrid NRPS/PKS cluster elicits induction

Because only 12-B09 and FS-144 elicited lysis and prophage induction among the five *V. ordalii* strains in our collection (fig. 1A-C, fig. S4A), we were able to identify one putative biosynthetic gene cluster (BGC) specific to the inducing strains using bacterial antiSMASH (v7.0)(*23*) (fig. S4C). This BGC constitutes a hybrid nonribosomal peptide synthetase (NRPS)/polyketide synthase (PKS) gene cluster - hereon the *ord* cluster (fig. 2A). In total, the *ord* cluster comprises genes encoding a 4-phosphopantetheinyl transferase (*ordA*), two nonribosomal peptide synthetases (NRPSs; *ordB, ordD*), a polyketide synthase (PKS; *ordC*), a peptide deformylase (*ordE*), and a SAM-dependent methyltransferase (*ordF*) (fig. 2A). We leveraged the NRPS and PKS domain and substrate predictions to propose structural elements and a putative biosynthetic pathway for the *ord* metabolite using a combination of bioinformatic tools (antiSMASH, PRISM, and University of Maryland NRPS predictor, see Materials and Methods)(*23–28*). These analyses suggested that the metabolite is a nonribosomal peptide-polyketide product beginning with two L-valine residues loaded by OrdD, followed by an L-leucine or L-isoleucine residue incorporated by OrdB, and ending with the addition of one unit of malonyl-CoA by OrdC (fig. S5A, tables S1-3). The ketoreductase and terminal reductase domains of OrdC indicate the likely reduction of the β-ketone of the initial PKS elongation product to a secondary alcohol and generation of an aldehyde or alcohol at the terminal carbon atom of the natural product. Sequence and structural analysis of the OrdC thiolation and reductase domains revealed conserved catalytic residues suggesting functional domains in contrast to the incomplete domains predicted by antiSMASH(*23*) (fig. S5B-E). Finally, interrogation of OrdF’s sequence resemblance to known methyltransferases suggested its role as an *O*-methyltransferase, rather than an *N*-methyltransferase; however, bioinformatic analyses alone could not definitively predict the chemoselectivity of this enzyme. In contrast, while OrdE was annotated as a peptide deformylase, its role in biosynthesis could not be immediately linked. Interestingly, a previous study identified this same BGC in 12-B09 as responsible for the ability to kill a broad diversity of *Vibrio* in a strikingly similar skewed distribution to our screen (<5% of strains inhibited >25% of their tested strains)(*16*). Thus, we hypothesized that the *ord* cluster encodes the prophage inducer.

**Fig. 2.**
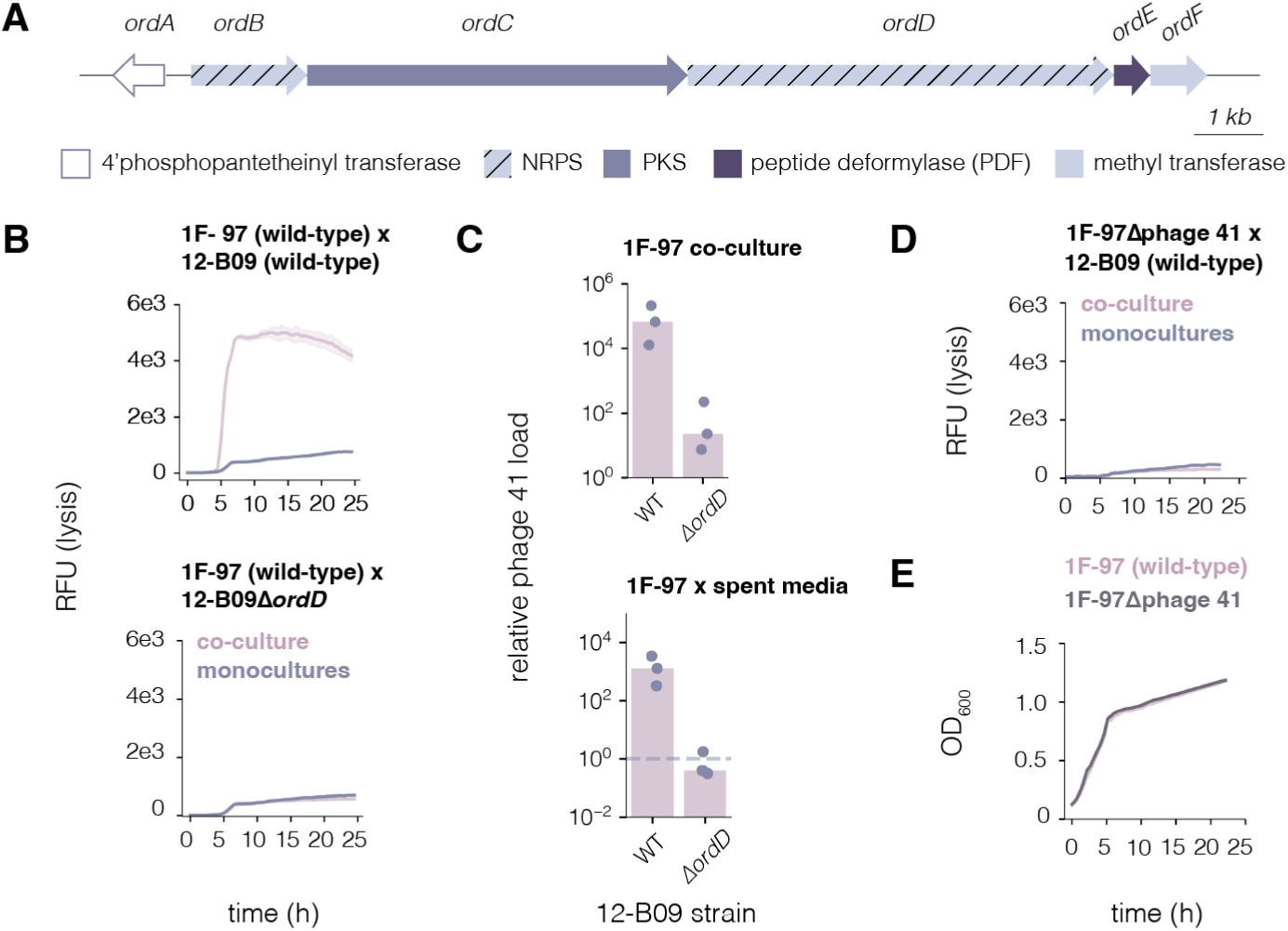
*V. ordalii* isolates elicit phage 41 induction using the *ord* NRPS/PKS metabolite. (**A**) The *ord* cluster, as predicted by antiSMASH (bacterial version 7.0(*23*)), is the only BGC unique to the prophage-inducing *V. ordalii* isolates 12-B09 and FS-144, but not found in the remaining *V. ordalii* in the collection (see fig. S4C). The putatively responsible metabolite appears to be synthesized from a hybrid nonribosomal peptide synthetase (NRPS)/polyketide synthase (PKS) cluster. (**B,C**) Comparisons between wild-type 12-B09 and 12-B09Δ*ordD* (null mutant disrupting *ord* cluster biosynthesis), demonstrating the change in the mutant to cause (**B**) lysis in a co-culture with 1F-97 and (**C**) phage 41 induction by spent media. (**D**) Lysis measurements of a co-culture between wild-type 12-B09 and an 1F-97 strain cured of phage 41. (**E**) Growth measurements to compare 1F-97Δphage 41 to wild-type. (B,D,E) Representative curve from three biological replicates, each with four technical replicates (mean with standard deviation). (C) Bars represent the median of three biological replicates (each with three technical qPCR replicates) shown with the individual biological replicates.

We confirmed that the *ord* cluster controls prophage activity by using a 12-B09Δ*ordD* mutant, where production of the metabolite is disrupted by deleting the central NRPS gene *ordD* (previously 12-B09 HW44 null mutant(*16*)). In contrast to wild-type 12-B09, 12-B09Δ*ordD* did not elicit lysis when co-cultured with 1F-97 (fig. 2B) and induction of phage 41 both in co-culture and by spent media declined severely (fig. 2C). Together, these data not only suggest that the *ord* cluster is primarily responsible for the induction of phage 41, but also that lysis by prophage induction appears to be the primary means for antagonism against 1F-97. To further assess whether lysis (proxy for antagonism) occurs exclusively due to prophage induction, we cured phage 41 from 1F-97. Similarly to the effects by 12-B09Δ*ordD*, co-culturing 1F-97Δphage 41 with wild-type 12-B09 results in a complete loss of lysis (fig. 2D). Furthermore, loss of phage 41 does not cause growth defects to 1F-97 in monoculture (fig. 2E), supporting the interpretation that 12-B09 antagonises 1F-97 exclusively through eliciting the induction of phage 41 with this hybrid NRPS/PKS product.

### Induction by peptide deformylase inhibition

We noticed that the *ord* cluster encodes a peptide deformylase (*ordE*) that did not have an apparent function in light of the proposed biosynthetic pathway (fig. S5A) - for example, the BGC does not also encode a formylase. Given that resistance genes often appear in BGCs, we hypothesized that the product of the *ord* cluster is a peptide deformylase inhibitor (PDI) of the highly conserved *N*-formylmethionine peptide deformylase (*def* gene or PDF product) and that OrdE was its resistance gene. In bacterial translation, formylated methionines are added as the first residue of every peptide, but their retention leads to the degradation of the nascent polypeptide(*29*). Thus, peptide deformylases must outcompete the degradation machinery to remove the formyl group soon after the nascent polypeptide is exposed at the exit tunnel of the ribosome. Furthermore, PDF is an essential gene, canonically encoded as one chromosomal copy upstream of the transformylase gene *fmt*(*29*). Given its importance for successful translation, inhibition of this process could plausibly serve as an SOS-independent stress signal prophages would be advantaged to detect and respond to. Notably, the previous study that had identified *V. ordalii* 12-B09 as a “superkiller” (capable of antagonising >25% of other *Vibrio* isolates) attributed its killing activity also to the *ord* locus(*16*). Without characterizing its mechanism, the killing was assumed to be by broad-spectrum antibiotic activity, potentially lysing cells by membrane disruption based on the biosynthetic gene cluster annotations(*16*). Here, we posit that the production of the *ord* metabolite could mediate killing by inhibiting the deformylation of nascent peptides, thus triggering induction in lysogens at otherwise sublethal concentrations to prophage-free cells. To that end, we first titrated actinonin - a commercial PDI(*30*), to observe whether it has activity against wild-type 1F-97 and whether the activity concentration differs from the concentration required to kill 1F-97Δphage 41. Lysis occurred in wild-type 1F-97 when treated with as low as 250 nM actinonin whereas lysis was only observed beginning at 2.5 µM in 1F-97Δphage 41 (fig. 3A). Furthermore, of the three prophages in 1F-97, only phage 41 was robustly induced by actinonin, optimally at 1 µM (fig. 3B). Together with the presence of *ordE* in the BGC, the shared capability to specifically induce phage 41 suggests that the *ord* metabolite acts by a similar mechanism to actinonin, that is PDF inhibition.

**Fig. 3.**
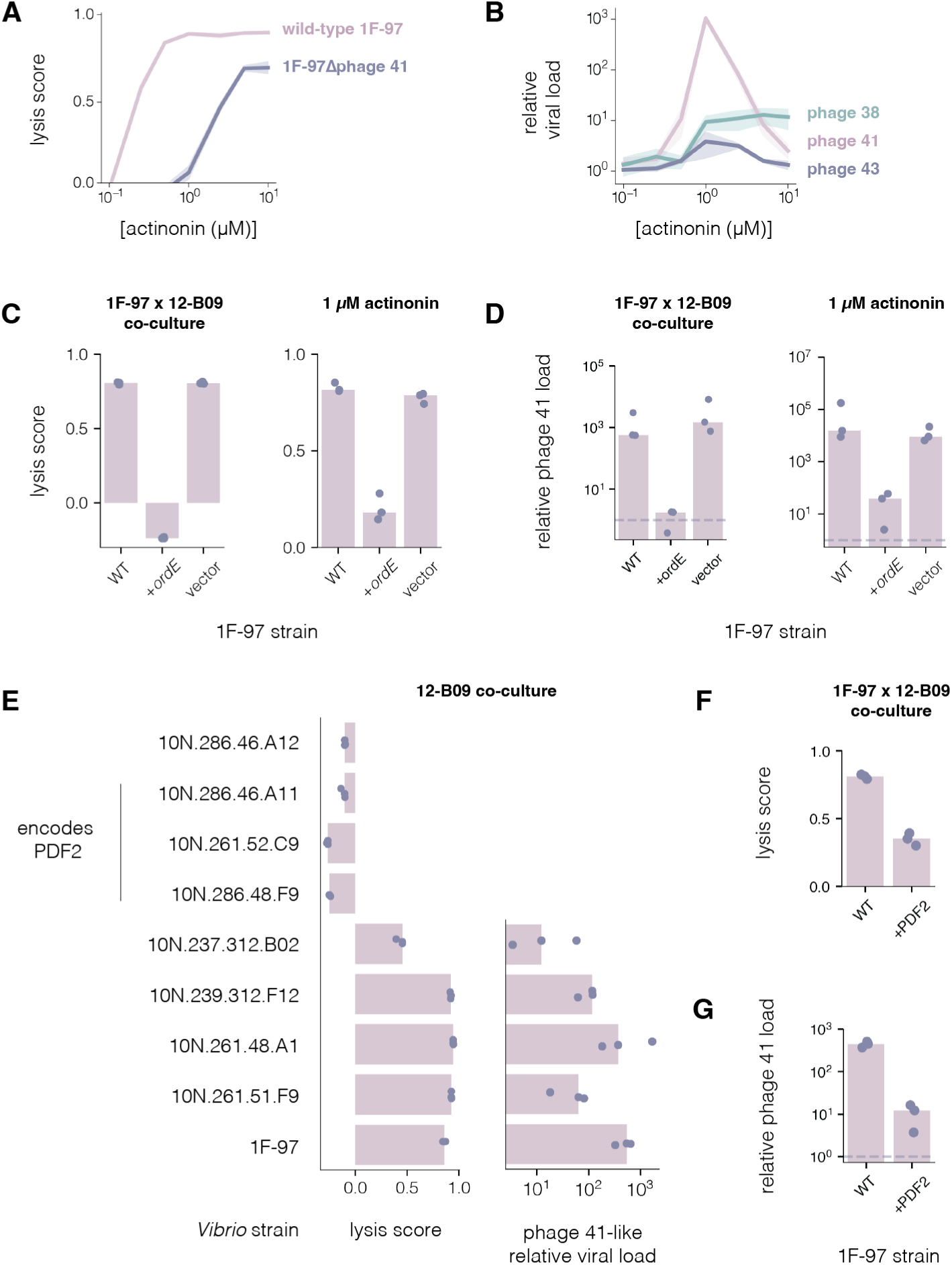
Phage 41 is induced upon peptide deformylase inhibition. (**A,B**). Titration of actinonin against 1F-97 and the respective (**A**) lysis scores (both wild-type or Δphage 41) and (**B**) relative viral load for phages 38, 41, and 43 from wild-type 1F-97. (**C**) Comparison of lysis scores between 1F-97 strains (wild-type, heterologously expressing *ordE* by its native promoter, or empty vector). The three strains were grown in co-culture with *V. ordalii* 12-B09 or treated with 1 µM actinonin during logarithmic phase. (**D**) A comparison of relative viral loads of phage 41 between the same 1F-97 strains seen in panel c. (**E**) Panel of *Vibrio* hosts carrying prophages resembling phage 41 with the respective lysis scores and relative viral loads when co-cultured with 12-B09, all at starting inocula of OD_600_ 0.05. A line denotes which hosts encode a second peptide deformylase (PDF2), in addition to the canonical PDF. (**F**) Aggregated lysis scores and (**G**) relative viral loads when PDF2 from 10N.286.48.F9 is heterologously expressed in 1F-97 by its native promoter. (A,C,E,F) Lysis scores generated from the mean of three biological replicates and the standard deviation from four technical replicates. (B,D,E,G) Bars represent the median of three biological replicates, each as the mean of four technical replicates shown (B) with standard deviation or (D,E,G) as individual points. (D,G) The dashed line represents no difference (1X) from the untreated 1F-97 culture.

Next, to evaluate if *ordE* is indeed the resistance gene of the *ord* cluster, we assessed whether heterologously expressing *ordE* conferred resistance against peptide deformylase inhibitors. Upon its expression in 1F-97 from its native promoter, we did not observe growth defects in the absence of induction (fig. S6). When co-cultured with 12-B09 or treated with 1 µM actinonin, 1F-97 no longer exhibited lysis while expressing *ordE* (fig. 3C). Consistently, phage 41 was also no longer induced in 1F-97 (fig. 3D). Altogether, we demonstrated that *ordE* sufficiently rescues 1F-97 from induction by both the *ord* metabolite and actinonin, thus concluding that *ordE* is encoded in the *ord* cluster as its resistance gene.

To test how broadly the *ord* metabolite can lead to prophage induction in additional strains, we bioinformatically identified prophages similar to phage 41, yet observed that some hosts were resistant to experimental induction in co-culture. Among the ten additional prophages found in nine *Vibrio* hosts, 12-B09 only displayed activity against four additional strains (fig. 3E, Materials and Methods). Phage 41 and all ten relatives form their own family in the *Caudoviricetes* together with other Vibriophages. In this family, phage 3 of 10N.286.46.A11 forms its own new genus, but the remaining nine belong to another new genus comprising eight new species (the relative in 10N.286.46.A12 and phage 1 of 10N.286.46.A11 were deemed the same) (fig. S7). Given the high degree of genetic similarity and synteny between these phages, but the absence of uniquely shared genes by the non-sensitive phages (fig. S8A), we speculated either that resistance was a host property rather than encoded in the phages, or that the prophages had become cryptic (i.e. lost induction capacity). Pointing to resistance being a host property, given how the *ord* cluster encodes its own peptide deformylase OrdE, we discovered that three out of four resistant hosts contained a second PDF gene (PDF2), while inducible strains only carried the canonical PDF (PDF1) (fig. 3E). PDF1 and 2 represent divergent protein clusters (up to 50.9% amino acid identity), while being highly similar within a cluster (97-100% identity) (fig. S8B, data file S2). Moreover, OrdE only shared up to 48.5% and 61.5% identity with PDF1 and 2, respectively. Despite the low sequence similarity, all three PDF clusters maintain the three conserved motifs around the catalytic site (G43XGXAAXQ, E88GCLS, and H132EXXH)(*31*) and share structural similarities (fig. S8C), suggesting that they are functional as PDFs but may have adapted to different constraints. Therefore, we hypothesized that the resistant strains horizontally acquired a second copy of PDF to protect against such antagonistic interactions.

Finally, to test whether PDF2 confers resistance against the *ord* metabolite, we heterologously expressed PDF2 (encoded by *Vibrio lentus* 10N.286.48.F9) in 1F-97. Similarly to OrdE, expressing PDF2 from its native promoter reduced lysis and phage 41 induction (fig. 3F-G), but unlike OrdE, it did not protect 1F-97 from 1 µM actinonin (fig. S8D). However, 10N.286.48.F9 itself is not resistant to 1 µM actinonin either (fig. S8E-F), suggesting that PDF2 may have diverged to specifically prevent binding to the *ord* metabolite (and related compounds), while OrdE became more generally resistant (preventing the binding of both the *ord* metabolite and actinonin). Taken together, these data suggest that the *ord* metabolite triggers prophage induction at sublethal concentrations as a PDI, and that natural isolates have evolved resistance against the *ord* metabolite through alternative PDF2 genes.

### Prophage induction as competitive interference

The presence of resistance factors against the *ord* metabolite in natural isolates (PDF2) suggests that similar interactions occur in the wild and impact population dynamics. We therefore asked how prophage induction affects the coexistence of the 12-B09 producer and 1F-97 lysogen by comparing the growth of each in mono- and co-culture using isolate-specific qPCR. If interactions were solely determined by competition for resources, we would expect 1F-97 to have a fitness advantage given its faster growth rate in monoculture compared to 12-B09 (growth rates of 0.9 h^-1^ and 0.8 h^-1^ respectively, and 12-B09 having a lag phase around 5 hours longer, see Materials and Methods). Indeed, both wild-type and 1F-97Δphage 41 strains dominated the culture when grown with 12-B09Δ*ordD* (fig. 4). Likewise, 1F-97Δphage 41 grew similarly when grown with wild-type 12-B09, likely due to the absence of prophage induction. However, when both wild-types competed, 1F-97 began to plateau around 12-h, presumably because prophage induction limited the abundance of 1F-97, while 12-B09 continued to grow. As further confirmation, we tested these four pairs by Burkholder assay and observed that only wild-type 12-B09 can cause clearance on wild-type 1F-97 (fig. S9A). These data validate the dependence of 12-B09 on induction by the *ord* metabolite for its ability to successfully compete with prophage-carrying strains. Finally, we asked whether the induction of phage 41 led to a subsequently detrimental infection of 12-B09. Upon screening 12-B09 colonies (n=44) for the presence of phage 41 by qPCR post co-culturing with 1F-97, the colonies resembled the negative control, suggesting that phage 41 does not lysogenize 12-B09 (fig. S9B). Together, these results describe how antagonism driven by prophage induction may serve as a successful form of interference competition.

**Fig. 4.**
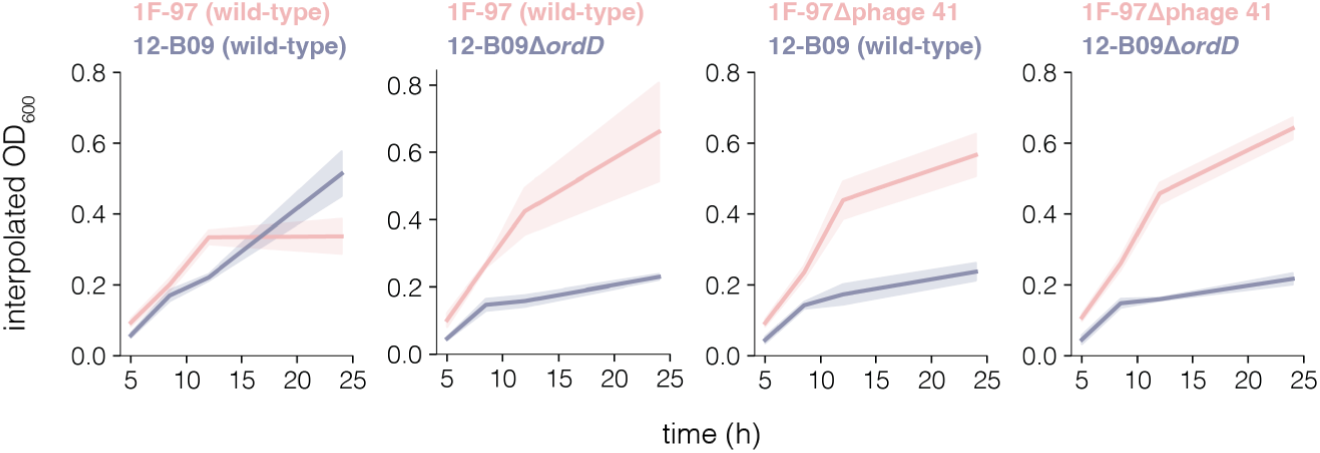
Prophage induction enables the coexistence of 12-B09 with 1F-97. Growth curves of each isolate from co-cultures where OD_600_ was interpolated by qPCR using a standard curve of known OD_600_ values and primers specific to the respective species (shared by strains). Relative growth of 12-B09 and 1F-97 were compared between all four combinations of pairwise co-cultures comprising 12-B09 (wild-type or Δ*ordD* in indigo) and 1F-97 (wild-type or Δphage 41 in orange) to assess fitness changes between conditions. Time points were taken after 5, 8.5, 12, and 24 hours and are represented as the mean of three biological replicates (each with three technical qPCR replicates aggregated to its mean) and standard deviation.

## Discussion

While genome mining discovers an abundance of biosynthetic gene clusters, bioinformatic analyses alone typically cannot accurately ascribe the mechanisms of actions to their products. Often, and unsurprisingly given strong prior expectations, these natural products are classified solely as antibiotics. For example, killing by the *ord* metabolite was previously assumed to be as a broad-spectrum antibiotic. However, despite no semblance to known inducing agents, we demonstrate that the *ord* metabolite can directly induce prophages. Moreover, this metabolite links translational stress to prophage induction which, to our knowledge, has not been previously described. Together with autoinducers, there now appears to be a diversity of SOS-independent prophage inducers(*13*, *14*), suggesting that additional classes and mechanisms of inducing compounds likely remain to be discovered.

By further broadening the types of compounds capable of inducing prophages, we begin to see differences in responses by prophages. Similarly discussed in other studies about polylysogenic strains(*22*), the selective induction by the *ord* metabolite suggests that prophages respond to multiple, distinct, induction cues. While the widespread SOS-responsivity may result in a race for host resources between competing prophages, uniquely responding to a specific cue, such as peptide deformylase inhibition, provides additional selective advantage under certain environmental conditions, which may partially explain how the coexistence of multiple prophages can be evolutionarily maintained in a single host.

Notably, because the mechanism of action was unknown, the previous antagonism study not only mistook the *ord* metabolite solely as an antibiotic, but also did not identify the resistance gene for the *ord* cluster with traditional genetic methods despite how immunity is typically encoded within a BGC(*16*). Studies on the antibiotic activity of actinonin and similar peptide deformylase inhibitors have revealed that resistance develops through spontaneous loss-of-function mutations in the *fmt* gene, rather than in the *def* gene(*32*, *33*). While lack of initial formylation to methionine shares similar genetic logic to redundant function by a second PDF, mutants lacking a functional methionyl-tRNA formyltransferase tend to be fairly sick, explaining why the primary method employed, transposon-mutagenesis, had struggled to characterize the mechanism in the past study(*16*), (*32*, *33*). Moreover, recent work elsewhere revealed that nearly 50% of available bacterial genomes encode additional predicted PDFs, often within the mobilome, highlighting a potentially widespread yet previously unrecognized resistance mechanism(*34*). Furthermore, a similar BGC, also thought to produce a peptide deformylase inhibitor (PDI) and encode a PDF for self-immunity, has been recently identified in *Xenorhabdus* spp. and other *Gammaproteobacteria*(*35*). The abundance of PDF enzymes and distribution of PDIs not only suggests the underappreciated ecological significance and surprising prevalence of PDIs, but also underscores the importance of disentangling the role of phages in shaping the selective pressures that have driven such remarkable diversification and maintenance of the extensive repertoire of PDF enzymes.

Finally, broadening the classes of prophage inducers also has translational implications and supports new antibiotic therapeutic hypotheses. Peptide deformylase has been an unattractive target for drug development due to the limited range of potent inhibitor scaffolds discovered, high rates of resistance from spontaneous *fmt* mutants, and off-target activities(*36–38*) - at least, when PDIs are used as conventional antibiotics at high concentrations against peptide deformylase as an essential gene target. In contrast, our work demonstrated how this antibiotic instead unconventionally and sufficiently kills at low concentrations in certain lysogenic strains. Furthermore, it has already been posited that many antibiotics may exhibit non-antibiotic functions at their natural production level, especially when the high concentrations necessary for antibiotic activity are rarely achieved under physiological conditions(*1*). Combined with the unknown bioactivity of most identified BGCs, future efforts in the discovery of clinically-relevant antibiotics may benefit from considering the potential of compounds as dose-dependent and multimodal with alternative mechanisms of action like prophage induction.

## Acknowledgements

We thank F. Le Roux (University of Montréal) for assistance with *Vibrio* genetics. We also thank S. Pollak (University of Vienna), members of the Blainey Lab (MIT/Broad Institute of MIT and Harvard), in particular A. Le and F. Keer, and members of the Polz Lab (University of Vienna) for their scientific discussion or input on the manuscript.

## Funding Sources

Advanced Research Projects Agency for Health (ARPA-H) (1AY2AX000005-01) (J.C., P.C.B.) Natural Sciences and Engineering Research Council of Canada PGS-D Fellowship (567809 - 2022) (J.C.)

National Institute of Health (Grant 5R01GM132564-04) (K.P., A.M.C., E.P.B.)

E.P.B. is a Howard Hughes Medical Institute Investigator

National Science Foundation Graduate Research Fellowship (DGE 2140743) (A.M.C.)

Simons Foundation (Life Sciences Project Award-572792) (K.S., M.W., J.B., A.L., N.B., M.F.P.) Austrian Science Fund (FWF) [doi.org/10.55776/COE7] (K.S., M.W., J.B., A.L., N.B., M.F.P.) EMBO (ALTF-584-2022) (J.B.)

HFSP (LT0033/2023-L) (J.B.)

Marie Curie Postdoctoral Fellowship (Project 101063227 PHAGECOUNTER) (A.L.) 2021 Schmidt Science Fellowship (F.A.H.)

## Author Contributions

Conceptualization: J.C. with input from M.F.P.

Methodology, data curation, and formal analysis: J.C. with input from P.C.B.

Investigation: J.C., K.P., A.M.C., M.W.

Validation: J.C.

Software: J.C., K.S., N.B.

Resources: J.C., J.B., A.L., F.A.H.

Funding acquisition: M.F.P., E.P.B., P.C.B.

Visualization: J.C., K.P., N.B.

Writing – original draft: J.C., K.P., A.M.C.

Writing – review & editing: J.C., K.P., M.F.P., E.P.B., P.C.B. with contributions from all coauthors.

## Data and materials availability

The lysis screen datasets (both kChip and microtitre plate), qPCR prophage induction screen dataset, and fastq files (screen dataset and individual cultures, including information about the prophages detected), have been deposited onto Zenodo: https://doi.org/10.5281/zenodo.15131811. New genomes will be uploaded onto NCBI and accessions will be available upon publication. Plasmid maps will be available upon publication. Custom code used for kChip analysis, microtitre plate screen analysis, and prophage induction detection from Illumina short reads can be found on GitHub: https://github.com/chenjul/ord_induction. Further information and requests for resources should be directed to and will be fulfilled by Paul C. Blainey or Martin F. Polz.

## Supplementary Materials

Materials and Methods

Figs. S1 to S10

Tables S1 to S3

References (37-81)

Data S1 to S2

## Materials and Methods

### Bacterial strains and growth conditions

A list of strains and media used in this study are found in data file S2, including media additives and selection antibiotics(*39*). *E. coli, P. aeruginosa,* and *A. baumanii* strains were grown in Lysogeny Broth (LB; BD-Difco) and *S. aureus* with Tryptic Soy Broth (TSB; BD-Difco) with aeration or on LB/TSB agar at 37°C. On agar plates, *Vibrio* strains were grown on Marine Broth 2216 (MB; BD-Difco). As liquid cultures, *Vibrio* strains were grown in MB or a defined, minimal media broth (“M9G+”). *Vibrio* strains were grown at ambient temperature (for kChip/agar plates) or 22°C at 250 RPM (aerated cultures), unless otherwise specified. Microtitre plates were incubated in a BioTek Cytation 5 plate reader at 22°C, cooled with a Peltier unit, with intermittent 5-s shaking every 30 minutes to better resemble growth conditions in the kChip. *E. coli*, *A. baumanii, P. aeruginosa,* and *S. aureus* were grown in slightly adjusted M9 minimal media for SYTOX Green lysis experiments. We used M9 minimal media instead of TSB when measuring the interaction between *P. aeruginosa* and *S. aureus* because of its lower autofluorescence.

For the kChip and microtitre plate screening, strains were revived by pin-replication from 96W glycerol stocks into 200-µL of MB in 0.8-mL 96W deep-well plates and grown for 24-h. To acclimate cultures in a more physiologically-relevant environment (e.g. nutrient-limited), MB cultures were then diluted 1:100 into 400-µL of M9G+ in 2-mL 96W deep-well plates (“pre-cultures”), grown for another 24-h. For all remaining, non-screening experiments, strains were revived as single colonies on MB agar, then acclimated as above if grown in M9G+. For pooled sequencing and qPCR follow-up, grown pre-cultures were diluted to the desired initial culture inocula at a final volume of 1.6-mL in M9G+, represented in eight wells for pooled sequencing (12.8-mL per co-culture in a pool, 1.6-mL per well in 2-mL deep-well plates) and one per biological replicate for qPCR (1.6-mL), grown for 24-h. Other qPCR experiments measuring relative viral load were typically grown as 400-µL in 2-mL deep-well plates. To achieve conditions more similar to the intermittent shaking in microtitre plate experiments, shaking was reduced to 200 RPM. For spent media sources, grown pre-cultures were diluted 1:100 into M9G+ once more and grown with aeration at 22°C for 24-hours. The supernatants from the pelleted cultures were filtered through 0.22 µm PES filters (Corning SteriCups). To account for the depletion of nutrients, the spent media was supplemented with 20X solutions of select nutrients, diluted into the spent media to achieve a final minimum of 1X of M9 salts, Casamino acids, D-glucose, MgSO_4_, and CaCl_2_ (“supplemented spent media”). Finally, to measure relative abundance of strains in co-culture by qPCR, co-cultures were grown in 5-mL in 25-mL culture tubes. To calculate growth rate, an OD_600_ plate reader time courses (100-µL if 96W and 50-µL if 384W) were fit to an exponential curve (including 12-B09’s additional 5-h long lag phase despite media acclimation) using `scipy.optimize.curve_fit` on `K / (1 + np.exp(-r*(t - t_0)))`.

### Assays measuring interactions

To estimate bacterial cell lysis, cultures were incubated with a final concentration of 5 µM SYTOX Green. An empirical, error-adjusted “lysis score” was calculated using the area between the co-culture and the sum of the respective monocultures SYTOX Green fluorescence curves or between treated and untreated curves. Timepoint measurements were summarised as the mean fluorescence and error-adjusted with the standard deviation (subtracted from co-cultures, added to monocultures). The difference between the areas of the resulting curves was scaled to the maxima of the two curves to account for the different maxima of different co-cultures, producing absolute values (scores) between 0 and 1. Scores were positive if the co-culture displayed a greater signal than the sum of the monocultures, whereas the scores were negative if the co-culture displayed a weaker signal. In total, lysis scores were found between -1 to 1. This calculation can be found in the provided code. Fluorescence was measured with the BioTek Cytation 5 plate reader using the following settings: 493 nm excitation, 534 nm emission, BPF 20, top reads, and gain 50. Details about fluorescence reads by microscopy can be found under kChip Screening.

For validation of the effects seen using the SYTOX Green lysis assay, the Burkholder agar assay was performed by mixing top agar (0.3% agar) with 100-µL of the overnight indicator culture, then poured over the bottom agar (1.5% agar). Once the 2.5-mL of top agar fully solidified, 3-µL of the overnight interacting culture was spotted onto the top agar. Plates were dried and incubated upright. Images were taken with an iPhone 13. Protocol adapted from Burkholder, P.R *et al*(*40*).

### Genome sequencing and bacterial species tree

For isolates ZF-76, 10N.261.54.E10, 5S-268, and the mutants built in this study, genomic DNA was extracted from 2-mL overnight cultures (MB for *Vibrio* strains) using the Qiagen DNeasy Blood & Tissue Kit. Sequencing, assembly, and annotations were performed by SeqCenter LLC. (Illumina NovaSeq 6000 with 2×151-bp paired-end reads; Unicycler v0.4.8, QUAST v5.0.2, and Prokka v1.14.5 with default parameters)(*41–43*). Additional Nanopore sequencing was prepared for 1F-97, 12-B09, FS-144, FS-238, FF-93, FF-167, and FF-266 with genomic DNA extraction as above. Sequencing, assembly, and annotations was performed by SeqCenter LLC. (porechop v0.2.4 with default parameters; flye v2.9.2 with --asm-coverage 50 --genome-size 6000000 --nano-hq; circulator v1.5.5 with all; 6-hour timeout; Bakta v1.8.1 with default parameters and db version 5.0; and QUAST v5.2.0 with default parameters)(*42*, *44–46*). *Vibrio coralliilyticus* YB2 and its sequenced genome was gifted by the Cordero Lab (MIT), further annotated with Bakta (v1.7.0, default parameters, db-light)(*46*). Finally, all remaining genomes published with this work were sequenced from 1.2-mL MB cultures, prepared with the Nextera DNA Library Preparation Kit, ran on a Illumina HiSeq 2500 (2×101-bp paired-end reads), assembled with UniCycler(*41*), then annotated with Bakta (v1.7.0, default parameters, db-light)(*46*).

From the whole genome sequences, a bacterial species tree was constructed from concatenated ribosomal proteins using an adapted version of the Snakemake workflow RiboTree (https://github.com/philarevalo/RiboTree). Tool parameters were set to those of the workflow, except at alignment trimming where, instead, columns were removed if they contained greater than 90% gaps and at tree generation where rapid bootstrapping used 100 replicates.

### Genome mining and bioinformatic predictions

To survey the prophages found in the strain collection, non-redundant prophage sequences were predicted from whole bacterial genome sequences using VirSorter2 v2.24, DeepVirFinder v1.0, and VIBRANT v1.2.1 adapted from the Snakemake workflow “Virsearch” (https://github.com/alexmsalmeida/virsearch)(47–49), appending additional predictions by geNomad v1.5.1(*50*). In summary, all prediction tools used their default settings other than minimum contig lengths of 10,000 (Virsearch tools) or using the “conservative” flag (geNomad). DeepVirFinder predictions were further filtered for scores > 0.9 and p-values < 0.05. Then, phage predictions were filtered by the quality assessments summarized by CheckV v1.0.3 (end-to-end default)(*51*). Generally, we aimed to reduce false positives over false negatives, filtering by minimum contig and predicted phage sequence lengths of 10000, the greater presence of predicted viral genes than host genes, kmer frequencies ≤1 if applicable, and no “contig 1.5X longer than expected” warnings. Finally, filtered predictions were clustered with CD-HIT-EST (-c 0.99) to remove close-to identical phages(*52*). Given that the different prediction tools could define phages differently at their borders, CD-HIT-EST would miss some redundant predictions. Therefore, contigs with multiple predictions were analysed for similarity using MASH v2.3 (MASH distances ≤0.05 and p-values ≤1e-10, calculated from 25-mers and a sketch size of 400,000)(*53*) and BLASTn v2.13.0 (default parameters for each phage against a custom database of all predicted phages for the given host)(*54*). If predictions were deemed redundant, the shorter prediction was removed from the final set of predicted prophages.

In order to identify the metabolite responsible for induction, a comparative analysis was performed using the biosynthetic gene clusters predicted with antiSMASH 7.0(*23*). To further annotate a biosynthetic gene cluster, the 50-kbp region of the *Vibrio ordalii* 12-B09 genome hypothesized to encode the inducing agent (and contains deletion locus that resulted in loss of the prophage induction activity, as 12-B09*ΔordD*) was analyzed using antiSMASH 7.0(*23*), PRISM 4(*27*), and the University of Maryland PKS/NRPS analysis tool(*28*) in May 2024. The putative domain architectures and substrates identified by these tools (tables S1-3) were used to generate a predicted metabolite structure and to inform deconvolution of the major mass feature identified in the subsequent comparative metabolomics experiments. The NCBI Conserved Domain Database(*55*, *56*) and BLAST searches(*54*) in the UniProt and NCBI databases were used to inform the activity of methyltransferase OrdF.

To assess the divergence between peptide deformylases, peptide deformylases were identified from Bakta annotations (see above). A gene tree was generated using MUSCLE v5.3(*57*) alignment of all peptide deformylase amino acid sequences, FastTree v2.1.11(*58*) with the WAG+CAT model, and visualized with MEGA X v10.1(*59*). The structures of the peptide deformylases were predicted using AlphaFold 3(*60*) to assess structural conservation.

### Similar phage identification and taxonomic classification

To search for phage similar to phage 41, phages were first identified among publicly available genomes by searching for conserved neighbourhoods for various genes from phage 41 using JGI’s Integrated Microbial Genomes & Microbiomes. After confirming experimentally which attainable genomes displayed lysis and induction (see “Assays measuring interactions” and “qPCR to measure relative viral load or interpolate growth”), we used phage 41 and similar phages from genomes 10N.261.51.F9, 10N.261.48.A1, and 10N.286.49.C5 to further search for phages against recently published genomes(*19*). Notably, 10N.286.49.C5 demonstrates atypically high lysis in monoculture, which obscures relative measurements and was thus not considered further in other analyses, but was kept to search for phages (fig. S10). To choose marker genes, ORFs were first re-identified with prodigal v2.6.3(*61*) from the tested phage dataset, subsequently constructing a similarity matrix of the proteins using blastp v2.13.0(*54*) (-eval 0.00001). Proteins were then clustered with mcl v22.282 (--stream-mirror --stream-neg-log10 -stream-tf ‘ceil(200)’). The most conserved protein clusters were chosen as marker genes for the search, which were annotated as large terminase subunit, ParA, and major tail tube (accessions: WP_372435030, WP_016785612, WP_016785647). From the new set of genomes, flexible genomes were calculated using ppanggolin v1.2.105(*62*) (panrgp mode) for different *Vibrio* populations, where a population is assigned if it contains at least 5 genomes (see Methods in Steensen et al(*19*). ORF and protein identification was performed as above (adjusted -eval= 0.001 for blastp) with these new genomes to identify whether the genomes had any of the three chosen marker genes. Genomes were grouped by the presence of different combinations of the marker genes, viewed using CLINKER v0.0.29(*63*), and genomes with all three elements were then similarly tested experimentally by lysis and induction as above.

To taxonomically classify the identified prophages, we extracted relatives using the INPHARED database (Jan 2025). We predicted all proteins with prodigal v2.6.3(*61*) using the meta procedure and ran a blastp v2.15.0(*54*) (bit score >50,e-value <0.00001) against the INPHARED(*64*) database. We filtered for relatives sharing at least 15% of the query phage genes. To determine family level clusters, VirClust(*65*) (standard parameters, 1000 bootstraps) was run and all phages falling in the same cluster with the identified prophages were further classified in viral genera and species using VIRIDIC(*66*) (standard parameters). In addition, a subset of phages representing all family-level clades was used for a VICTOR(*67*) analysis (“amino acid” and “d6” intergenomic distance formula).

### kChip screening

The kChip screening method has been described previously(*20*, *21*, *68*); the adapted method is summarised as follows (also described in fig. S1A). First, kChips were fabricated with microwell diameters of 143-µm and heights of 135-µm, fitting maximally 124,260 microwells in a single chip. Pre-cultures were diluted in M9G+ at low and high OD_600_ (0.02 and 0.1, twice the final inocula for t_0_ prior to droplet merging), supplemented with 0.05% (w/v) bovine serum albumin (SeraCare) which helps retain hydrophobic small molecules in droplets (e.g. barcoding dyes). The volumes of pre-culture used for these dilutions were determined by the OD_600_ measurement of 1F-97 and applied uniformly to all cultures. A final concentration of 5-µM SYTOX Green (Invitrogen) and 20-µM of the respective dye barcode (a unique ratio of Alexa Fluor 555, Alexa Fluor 594, and Alexa Fluor 647; Invitrogen #A33080, #A33082, #A33084) were added to the cultures. Cultures were dropletized using the BioRad QX200 Droplet Generator in a fluorocarbon oil (3M Novec 7500 with 2% 008-fluorosurfactant, RAN Biotechnologies). Emulsifications for each input were combined at equal volume and 800-µL of a uniform mixture was loaded onto the kChip, sealed with an optically clear PCR film (Applied Biosystems), wetted with 7500 oil. To map strain members to each co-culture assay, the kChip was first imaged at 1X magnification to record the barcoding dyes using the Nikon Ti-2 inverted fluorescence microscope (equipped with a Lumencor Sola light-emitting diode illuminator, Iris 9 camera, Nikon 1X Plan Achromat Microscope objective, and the filter cubes: Semrock SpGold-B, Semrock 3FF03-575/25–25 + FF01-615/24–25, Semrock LF635-B, and Semrock GFP-1828A). Droplets were then merged within their microwells with brief exposure to a corona treater (4.5 MHz, 10,000-45,000 volts; Electro-Technic Products, Model BD-20), bringing cultures to their final inocula and initiating the co-culture. The kChip was then imaged, again with all four channels, every 30 minutes for 24 hours (1X magnification). An all-by-all pilot kChip experiment (included in total dataset) involving 32 of the 110 *Vibrio* collection was first performed and used to select a positive control, which coincidentally was the 1F-97 x 12-B09 co-culture (0.05 x 0.05). Thus this pair was represented on and used for quality assurance for every kChip included in the screen dataset.

### kChip data analysis

Image analysis was performed using adapted, but previously published custom Python scripts(*68*). In summary, the combinations found in each microwell were decoded from the premerge images through the positioning of unique ratios of fluorescent barcode dyes for each input. This positioning was then mapped onto the respective merged images to associate the fluorescent lysis signal with the culture.

To estimate lysis from the kChip dataset, the median fluorescence was taken for each combination at each time point to produce empirical lysis scores (SYTOX Green lysis described above in “Assays measuring interactions” and fig. S1D). To estimate the standard error, we computed a distribution for each combination by bootstrapping (1,000 iterations) from all microwell technical replicates. We then calculated the standard error per time point using the standard deviation of the distribution. Our empirical lysis score was then error-adjusted by accounting the standard errors of the respective curves (i.e. subtracted the error from the co-culture median fluorescence or added the error to the respective monoculture median fluorescence, then re-calculated the AUCs/lysis score). Using the 1,000 iterations for each component in calculating a lysis score, we also computed a distribution of 1,000 lysis scores for each combination. Here, an estimated lysis score was derived from the mean of this distribution. From this point, all chips were processed as a single batch. To determine the significance of each score, we first standardized the estimated lysis score with its standard error (standard deviation of the score distribution) as 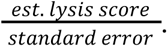 Benjamini-Hochberg-corrected p-values (q-values) were then calculated from these standardized scores using a nonparametric right-tailed test against the bootstrapped (10,000 iterations) null distribution which comprises the standardized scores of all “self co-cultures” (e.g. strain A_low_/strain A_high_). We reasoned that these cultures would not display antagonism while also better representing the resource use from the range of inocula in the dataset. In total, a combination would pass if the error-adjusted empirical lysis score ≥ 0.1, the FDR-corrected p-value ≤ 0.05, and the combination passed on at least two kChips. Finally, hits were binarized where a unique pair is called if at least one of the four ratios (combinations) passed the selection criteria. The best lysis score was that of the inocula ratio with the highest mean lysis score (calculated only from passing combinations).

To ensure the kChip dataset comprised only of chips with suitable effect sizes, chips were assessed by calculating Z-prime values, using 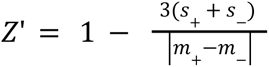and bootstrapping (1,000 iterations) similarly as above to compute standard errors (*s*). The maximum signal (*m_+_*) was represented by the chip’s 12-B09_high_/1F-97_high_ co-culture lysis score (AUCs calculated from median fluorescence); this co-culture was chosen as the positive control for its display of high lysis in the pilot with exceptions to chips missing one of the two inputs where 12-B09_high_/1F-97_low_ was used instead. The minimum signal (*m_-_*) was represented by the respective 1F-97 self-culture which displayed higher basal lysis levels than 12-B09. Chips passed if Z’ ≥ 0.7 and the error-adjusted empirical 12-B09/1F-97 lysis score ≥ 0.2 (at least twice that of the hit-calling threshold).

### Microtitre plate screening

M9G+ pre-cultures were grown and then diluted into their respective co-cultures or monocultures at final inocula of 0.01 and 0.05 in M9G+, unless otherwise specified, mixed with a final concentration of 5 µM SYTOX Green. Volumes of pre-cultures were calculated based on the 1F-97 culture’s OD_600_ measurement, as with kChip screening. Cultures were distributed as 4 x 50-µL replicates in 384W clear-bottom microtiter plates, sealed with a Breathe-Easy seal (Diversified Biotech), then grown in a BioTek Cytation 5 plate reader over a 22-h time course and read for both OD_600_ and fluorescence (see above for plate reader settings).

The empirical lysis score for each condition was calculated using the mean fluorescence for each timepoint. This score was then error-adjusted by the standard deviation - similarly to the kChip analysis. To determine the significance of each score, we performed a right-tailed test against the bootstrapped (10,000 iterations) null distribution which comprises the error-adjusted lysis scores of all “self co-cultures”. P-values were then corrected with the Benjamini-Hochberg Procedure. In total, a condition would pass if the error-adjusted empirical lysis score ≥0.1 and the FDR-corrected p-value ≤0.05. Downstream assays from this screen would go on to use initial inocula of each passing co-culture with the highest lysis score.

### Detection of prophage induction by NGS

Pooled cultures are described in “Bacterial strains and growth conditions” and cultures included in each pool are listed in data file S1. For pooled sequencing, cultures were pooled in equal volumes on ice after 24-h growth. Non-pooled cultures were grown in 25-mL. To obtain phage DNA, both pooled and unpooled cultures were extracted as follows. Cultures were centrifuged at 4°C for 15 min at 5250 g. To remove residual bacterial debris, the recovered supernatants were centrifuged again under the same conditions, then passed through 0.2 µm PES Stericup filters (Corning). Phage particles were further concentrated by PEG precipitation as follows. Filtered supernatants were incubated overnight (typically 16 - 18-h, up to 24-h) at 4°C at 4:1 of supernatant to 5X PEG solution (0.2 g/mL PEG-8000, 0.15 g/mL NaCl), then centrifuged at 15000 g for 30 min at 4°C. The recovered pellets were resuspended in TBS (8.8 µg/mL NaCl, 5 mM Tris-HCl, pH 7.5) to achieve the desired concentration (up to 100X), incubated on ice for 30 min to help dissolve the pellets as needed. To remove contaminating bacterial DNA, the phage resuspensions were treated with TURBO DNase I (Thermo Fisher Scientific) and RNAse A (Qiagen) at 3 U and 300 mg per 100 µL phage resuspension, respectively, for 2-h hours at 37°C, or until bacterial DNA was visibly absent via 1% agarose gel, then inactivated with 2 mM EDTA at ambient temperature for 10 min, followed by 75°C for 20 min. To release phage DNA from capsids, samples were treated 10:1 with 2.5% SDS (buffered with 0.5 M Tris-HCl, pH 9) at 75°C for 10 min, then brought to 55°C before digesting further with Proteinase K (Qiagen; at 0.2 mg/mL) at 55°C for 20 min, inactivating Proteinase K at 95°C for 10 min. DNA was extracted from cooled samples using 0.5X AMPure XP beads (Beckman) to avoid extracting small, residual bacterial fragments, then resuspended in ddH_2_O to achieve at least 500X concentration of the supernatant volume. Finally, libraries were prepared using the NEB Ultra FS II Library Prep Kit (typically input normalization to 0.2 ng/µL with 1 min fragmentation and 1/250 adaptor concentration, then amplified with 12 cycles) and loaded onto an Illumina NextSeq 500 (300 cycles), at approximately 0.5 M reads per condition in a pool and approximately 3 M reads per non-pooled samples.

After demultiplexing and conversion to fastq files (bcl2fastq2 v2.20.0), reads were mapped with Bowtie2 v2.3.4.3(*69*) against the respective host genome or a concatenated reference of all host genomes in a pool of samples. Briefly, the custom code called cases of prophage induction as follows. Sequencing depth was first calculated using samtools depth -a -H, including unpaired mates in the calculation. Next, we assumed that a phage is ≥10-kbp and reaches ≥10X coverage upon induction (median background-subtracted), thus a phage-containing contig would have a depth sum of ≥100,000 (the threshold is based on the sum basepair coverage of a 10-kbp region with exactly 10X coverage). To loosen the initial hit calling, contigs were filtered to have a depth sum of ≥50,000. To identify the borders of putative, enriched prophages, the contig was scanned for the first non-zero position whereby the following 50-bp contain fewer than 20% zero-coverage positions. To check whether it was a random patch of non-zero reads, the following 500-bp also required a minimum median coverage of 10X for the position to be considered an end coordinate. Finally, the spanning region was called an enriched prophage region if the median coverage had ≥ 10X coverage, its total length was ≥ 8,000-kbp (a looser cutoff than the assumed minimum 10-kbp phage genome length), and no more than 20% zero-coverage positions. However, to check whether a contig holds multiple phages and otherwise missed by the above cutoffs, we also manually checked the coverage using IGV v2.16.0(*70*) for contigs over 100-kbp with proposed start and stop coordinates.

### qPCR to measure relative viral load or interpolate growth

A list of all primers used in this study are found in data file S2. All qPCR experiments used KAPA HiFi HotStart 2X ReadyMix (Roche), SYBR Green (Invitrogen), and 1-µL of template per 25-µL reaction, measured using a CFX96 Real-Time PCR Detection System (Bio-Rad). The C_q_ values were generated using the CFX Maestro Software with the following settings: Regression (Cq Determination Mode) and Baseline Subtracted Curve Fit (Baseline Settings).

To detect prophage induction (relative viral load), the supernatants from grown, pelleted cultures were filtered with AcroPrep Advance 0.2-µm 96W PES filter plates, Corning 0.22-µm PES SteriCups, Whatman Cytiva GD/X 0.2-µm CA membrane syringe filters, or Corning Spin-X 0.22-µm CA tube filters, depending on volume and throughput. To remove bacterial DNA, the filtered supernatants were treated with TURBO DNase I (Thermo Fisher Scientific) at 2U per 10-µL of supernatant for 1-h, then inactivated with EDTA as in “Detection of prophage induction by NGS”, which subsequently contributes to releasing phage DNA from capsids for template. The relative viral load was measured by using the difference in quantification cycles (ΔC_q_) between the condition (co-culture, Mitomycin C, spent media etc.) and the uninduced monoculture of the host. Amplification was assumed to be 100% efficient for this estimate, thus the load was calculated as 2^ΔCq^. Purified genomic DNA was used as positive controls.

The abundance of unlabeled isolates grown together in co-culture was quantified using strain-specific primers for qPCR. To prepare the qPCR template at a specific timepoint, cultures were diluted 1:100 in ddH_2_O, then heat-killed at 100°C for 5 min. The OD_600_ for each isolate in a co-culture was estimated by interpolating from its respective standard curve (fitted as exponential decay using `scipy.optimize.curve_fit` on `m*np.exp(x*t)+b`) of known OD_600_ values and measured C_q_ values (from an overnight culture was diluted between a 0.1 to 1 OD_600_ range with a 0.1 step). Importantly, the standard curve was prepared using an overnight culture grown to late-exponential/early-stationary to capture a large OD_600_ range, while avoiding an overestimation of viable cells due to the presence of template DNA from increased cell death in stationary phase.

### Chemical prophage induction

For induction by Mitomycin C (MMC) or actinonin (Cayman Chemical Company), *Vibrio* cultures were grown from OD_600_ 0.01 to logarithmic phase (OD_600_ 0.2-0.35) with pre-culturing as described in “Bacterial strains and growth conditions”, then induced with a final concentration of 0.05 µg/mL MMC (or as specified for actinonin). For induction by spent media, targeted isolates were grown from OD_600_ 0.01 to approximately OD_600_=0.45-0.6 with pre-culturing as described above, and induced in equal parts with supplemented spent media. Induced cultures were grown for a total of 24-h from OD_600_ 0.01.

### Strain engineering

The list of constructs built for this study is found in the data file S2, with details about their respective primers, backbones, cloning strains, and method(*71*, *72*). All inserts were amplified using KAPA HiFi HotStart 2X ReadyMix (Roche) and backbones were treated with DpnI (NEB) as needed. Constructs were built using either Gibson Assembly (NEB 2X HiFi Assembly), replicated in *E. coli* Π3813 (replication strain) typically with a pSW7848T backbone or Golden Gate Assembly (NEBridge^®^ BsaI-HF^®^ v2), replicated in *E. coli* DATC (both replication and conjugative strain) typically with a pJB176 backbone. Colony PCRs were conducted with OneTaq (NEB). Purified constructs from *E. coli* Π3813 were then transformed into the conjugative *E. coli* β3914 (conjugative strain).

To cure all copies of plasmid-like phages, the putative origin of replication (a region flanking the *repA* gene) was inserted into pSW7848T for origin exclusion. Built constructs were introduced with the conjugation protocol below, then removed through the expression of *ccdB* toxin (counterselection). Once introduced into the *Vibrio* strain, the construct would either recombine or sequester host machinery away from the native phage origin when using a vector of much higher copy number, thus outcompeting an episomal phage. Single colonies were passaged daily on TSB-1+1% glucose+selective marker agar plates until intact phage was no longer detected by colony PCR (glucose to prevent *ccdB* expression). To remove the suicide vector, successful colonies were then replica-plated for counterselection onto both TSB-1+0.2% arabinose agar (to express the *ccdB* toxin if the vector remains present) and TSB-1+selective marker agar, grown overnight at ambient temperature. Colonies were passaged until they were able to grow on arabinose, but not on the antibiotic selective agar, then purified for single colonies. Finally, mutants were confirmed for their genotypes by Illumina short-read whole genome sequencing (SeqCenter) and mapped using Bowtie2 (v2.3.4.3)(*69*).

To disrupt a gene with a single recombination event, a region of the gene was inserted into pSW7848T with the *ccdB* gene, *araBAD* promoter, and *araC* removed. To disrupt *recA* in 1F-97, the *recA* gene was identified from the bakta(*46*) annotations and verified with HHpred(*73*) (web, using PHROGs_v4, PDB_mmCIF70_3_Jan, Pfam-A_v37, and UniProt-SwissProt-viral70_3_Nov_2021) annotations, also by its similar length to *E. coli*’s *recA* gene. Then, a 300-bp region of *recA* was inserted as above. Plasmids were introduced with the conjugation protocol below with no counterselection. Colonies were screened for successful recombination by colony PCR, then purified as single colonies onto MB+selective agar. As above, single colonies were passed daily until successful. For expression, the gene of interest was amplified and inserted in place of the CcdB toxin, introduced with the conjugation protocol below with no counterselection, and screened for successful conjugation via colony PCR.

### Vibrio conjugation

Overnight cultures of conjugative *E. coli* strain (LB) transformed with plasmids containing the desired construct and 1F-97 (MB) were centrifuged for 2 min at 8000 RPM at volumes to achieve a 3:1 donor-to-recipient ratio in 1-mL (strains used for specific constructs found in data file S2). Each pellet was washed separately with 500-µL of mating media (TSB-1+0.3 mM DAP, no antibiotic). After a second centrifugation, the *E. coli* pellet was resuspended in 500-µL of mating media, then transferred to the 1F-97 pellet. The combined cells were pelleted and resuspended in 30-µL of mating media. The resuspension was dropped directly onto TSB-1 agar plates with the respective additives and 1% D-glucose if the *ccdB* toxin is present on the backbone to prevent expression, then incubated upright for 24-h at 30°C to allow mating. Mating spots were then scraped and resuspended in 500-µL of SM buffer (data file S2). Serial dilutions were then plated on TSB-1+antibiotic (without DAP to ensure removal of the *E. coli* donor) and incubated for 36 - 48 hours at ambient temperature (∼22°C) until colonies appeared. Counterselection, if applicable, is described with the construct.

## Supplementary Figures

**Fig. S1.**
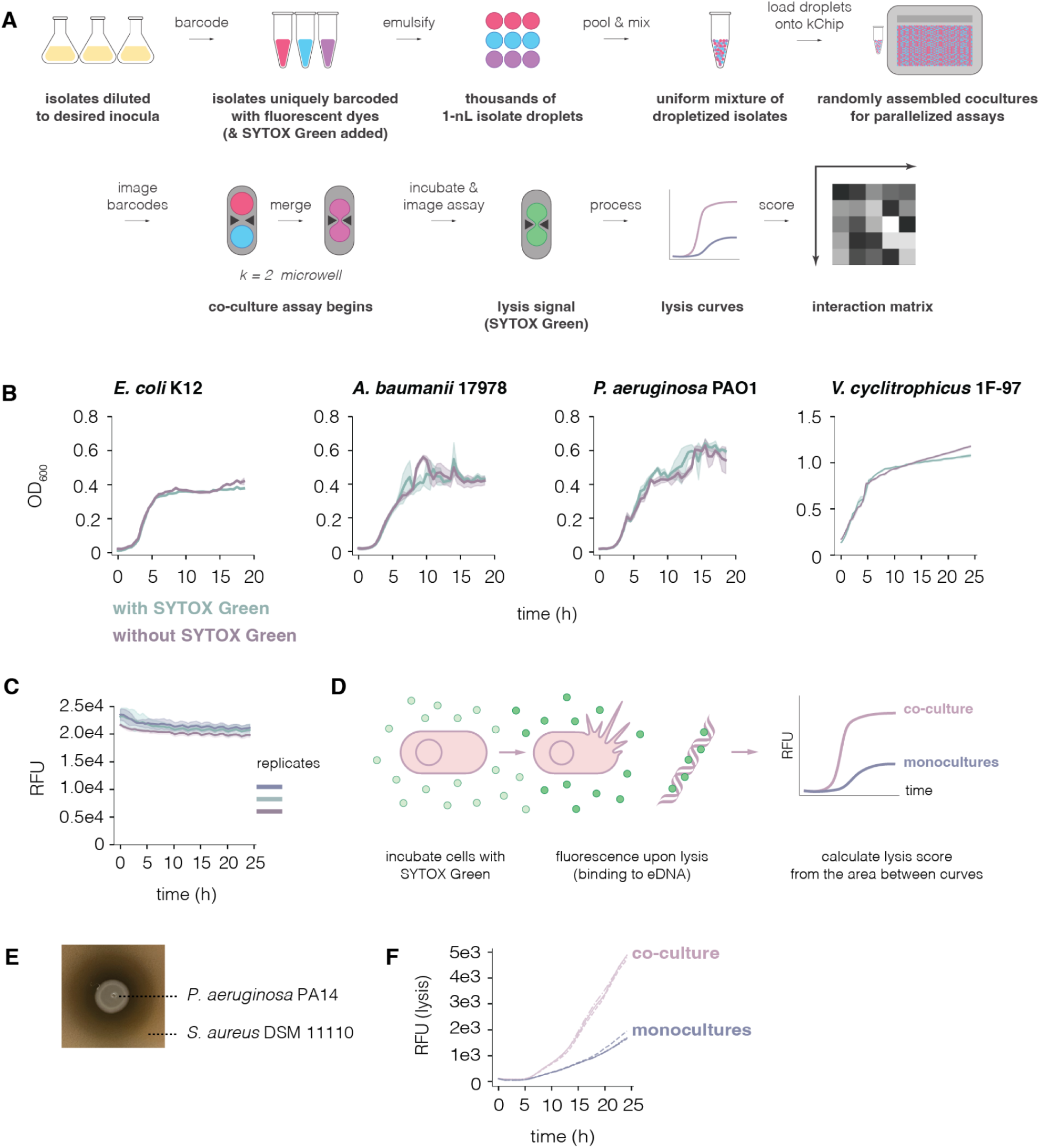
SYTOX Green combined with the kChip for high throughpout discovery of negative pairwise interactions leading to lysis. (**A**) Schematic of the kChip screen design to measure lysis-resulting interactions between isolates. (**B**) Growth curves of various species grown with (teal) and without (purple) 5 µM SYTOX Green in M9 minimal media. Representative curves from three biological replicates are shown in cultures growing for 18-h (*E. coli* K12*, A. baumanii* 17978*, P. aeruginosa* PAO1) or 24-h (*V. cyclitrophicus 1F-97*), performed in 384W microtitre plates, with four technical replicates per culture (as mean with standard deviation). (**C**) Fluorescence curves of 5 mg/100 µL of λ phage DNA with SYTOX Green to determine whether the dye is photobleached over the time courses used in this study. Three samples each with three technical replicates (as means with standard deviations) are represented in the plot. (**D**) Schematic describing how the cell-impermeable SYTOX Green can be used to detect cell lysis. Bacterial cultures are incubated with the dye at t_0_, displaying low fluorescence. Over time, fluorescence increases when the cell membrane is compromised or upon cell lysis (e.g. by antibiotics, prophage induction, etc.) as the dye can access and intercalate bacterial DNA. (**E,F**) A comparison between (**E**) the traditional Burkholder agar assay(*40*) and (**F**) the SYTOX Green lysis assay with *P. aeruginosa* PA14, which is known to antagonise *S. aureus* DSM 11110(*15*). (E) *P. aeruginosa* was spotted onto the targeted *S. aureus* on TSB agar, then incubated at 37°C for 18-h. The image is representative of three biological replicates, each with three technical replicates. (F) The lysis assay was performed with three biological replicates (each curve shown with different linestyles), each from the means of four technical replicates.

**Fig. S2.**
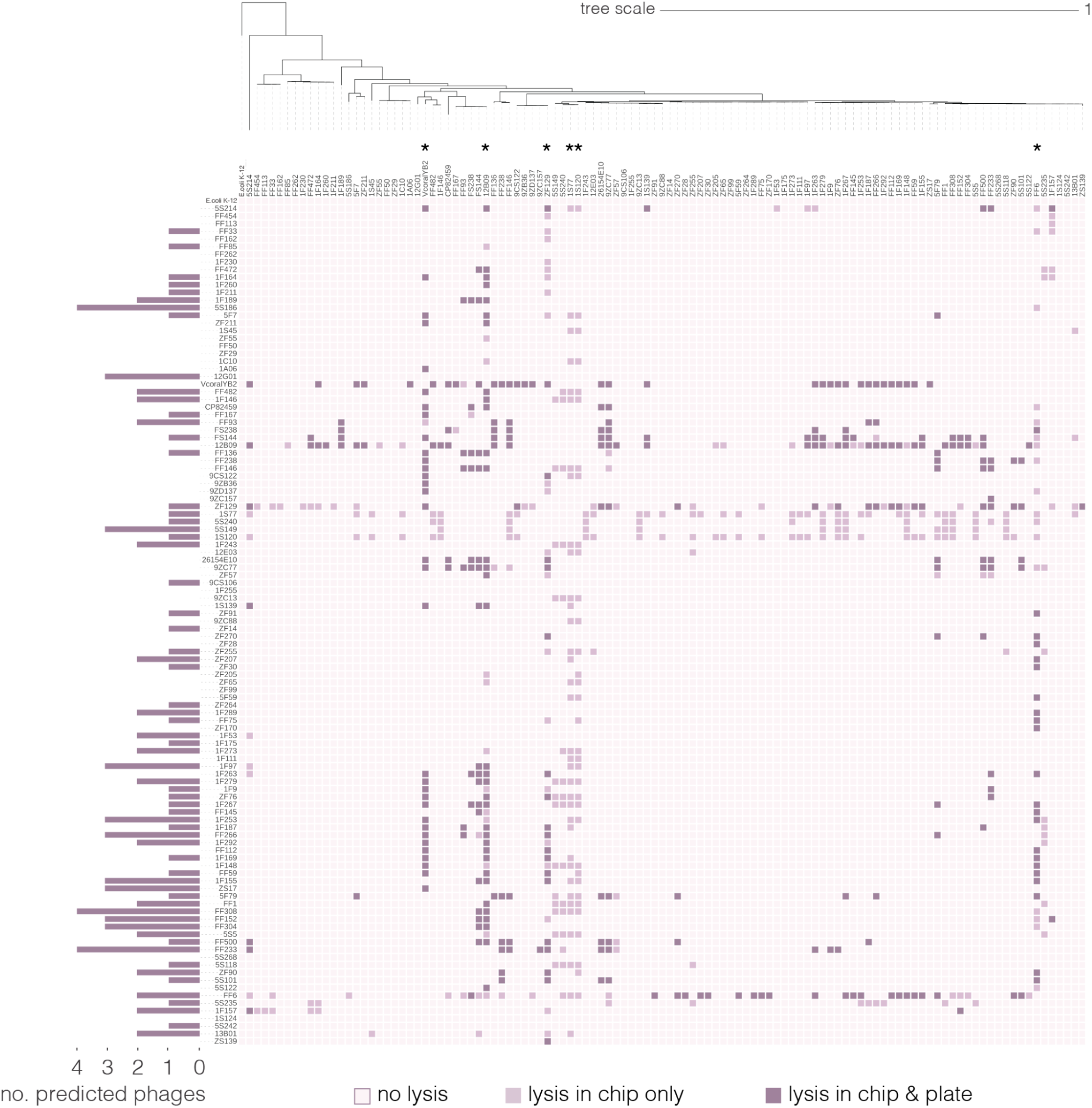
Negative interactions matrix resulting in lysis at different assay levels. Interaction matrix binarized based on the SYTOX Green lysis scores from an all-by-all pairwise screen from a collection of 110 natural *Vibrio* isolates, resulting in 344 (5.7%) of 5,995 unique isolate pairs that displayed lysis in the kChip. The screen comprised 5,995 unique isolate pairs, covered across 24,530 conditions (23,980 co-cultures and 550 monocultures with varying initial OD_600_ values of 0.01 or 0.05 per isolate) and an average of 14 replicates per condition. Isolate pairs passed if any of its conditions exhibited lysis. Initially scoring pairs were further downsized to the 172 (2.9%) interactions which were recapitulated in 384W microtitre plates. The darkest cells mark pairs that scored in both the kChip and microtitre plates, the lighter cells mark pairs that only scored in the kChip, and the lightest cells mark pairs that did not display lysis. The matrix is arranged by the collection’s phylogenetic tree on both axes, using *E. coli* K12 as the outgroup. Briefly, the tree was constructed using the ribosomal proteins from all isolates in the collection, aligned with MAFFT-L-INS-i(*74*) using their nucleotide sequences, and generated with RAxML(*75*) using the GTR model. The number of predicted phages per isolate are shown as a barplot along the left of the matrix. Isolates involved in lytic interactions with ≥25% of the collection are marked with an asterisk (*). The matrix was visualised using the iTOL annotation editor (v1.7) and webserver(*76*).

**Fig. S3.**
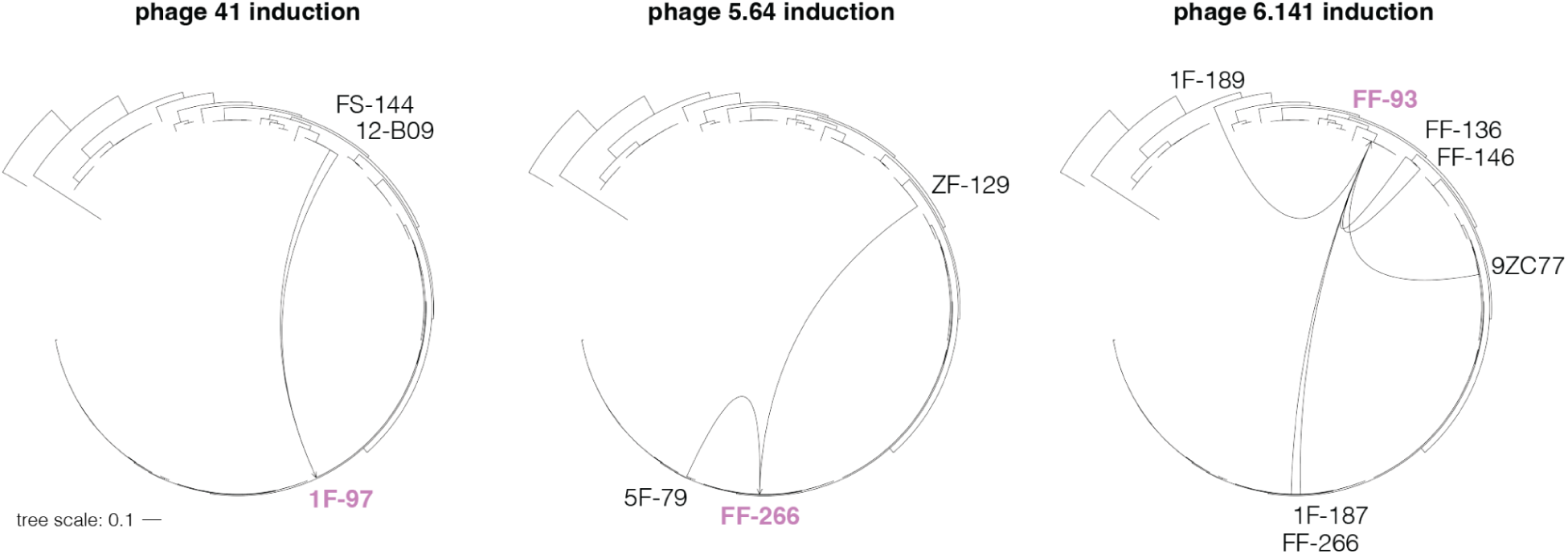
Phylogenetic relationships between inducing isolates and hosts of prophages inducible by co-culture. Ribosomal trees were constructed using MAFFT-L-INS-i(*74*) (nucleotide alignment) and RAxML(*75*) using the GTR model, then visualised using the iTOL annotation editor (v1.7) and webserver(*76*). The hosts of the co-culture-inducible phages 41, 5.64, and 6.141 are bolded and in purple (*V. cyclitrophicus* 1F-97, *V. tasmaniensis* FF-266, and *V. ordalii* FF-93 respectively). The edges connect the targeted host with *Vibrio* isolates that elicit induction (black labels).

**Fig. S4.**
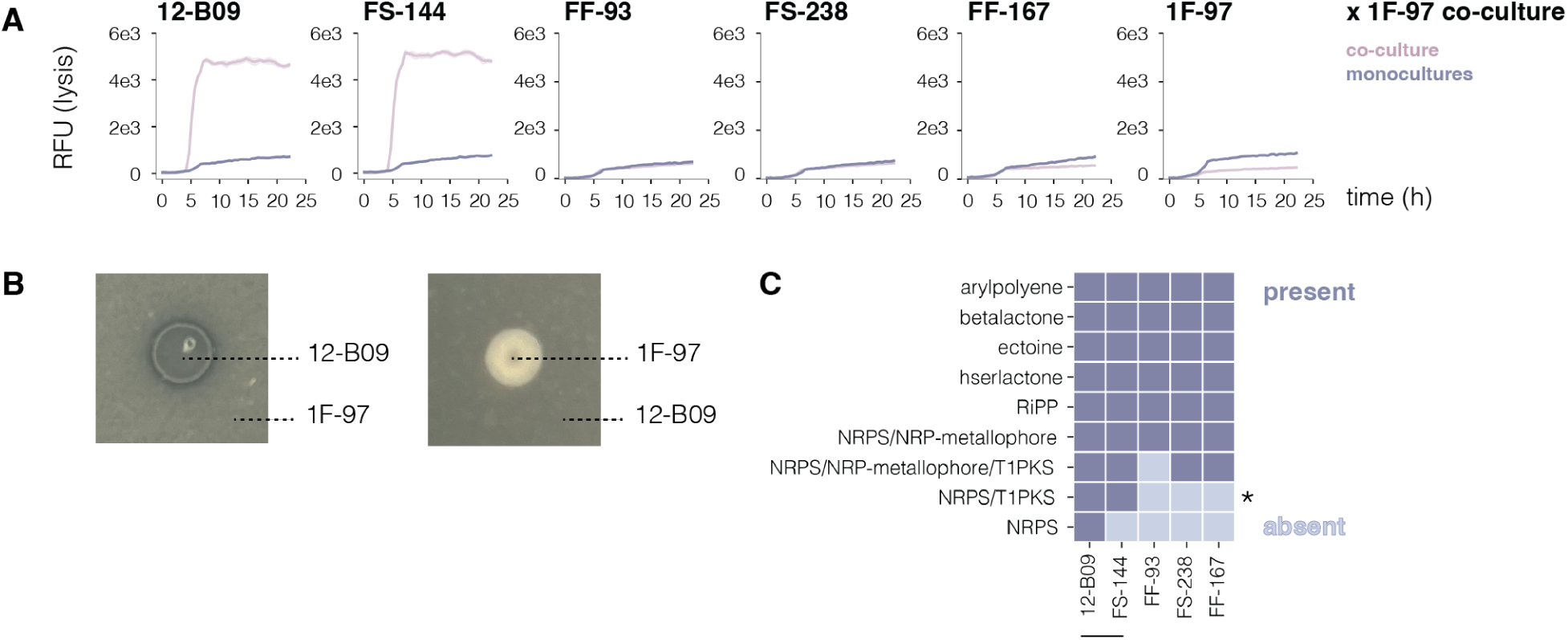
Shared NRPS/PKS cluster posited as responsible for lysis and non-spontaneous induction in 1F-97. (**A**) SYTOX Green lysis curves of co-cultures (pink) comprising 1F-97 with each of the *V. ordalii* isolates (12-B09, FS-144, FF-93, FS-238, FF167) or itself at starting inocula of OD_600_ 0.01 compared to the sum of their respective monocultures (indigo), associated with fig. 1A. Representative lysis curves from three biological replicates are shown from a 22-h time course performed in 384W microtitre plates, with four technical replicates per culture, represented as the mean with standard deviation. (**B**) Burkholder assay measuring the interaction between 12-B09 and 1F-97 in both directions. A dark ring around the potential antagonistic strain (drop spot) demonstrates a zone of clearance on the lawn of the other strain. Each image is representative of three biological replicates, each with three technical replicates. (**C**) Presence-absence matrix for predicted biosynthetic gene clusters in *V. ordalii* strains in the collection (antiSMASH v7.0(*23*)). Dark indicates the cluster is present, while light indicates it is absent from the strain; * marks the cluster proposed to be responsible for the induction of phage 41 as it is shared only by 12-B09 and FS-144 (marked by the line) - the only two isolates that elicit induction. Cluster class labels were assigned by antiSMASH using the MIBiG database.

**Fig. S5.**
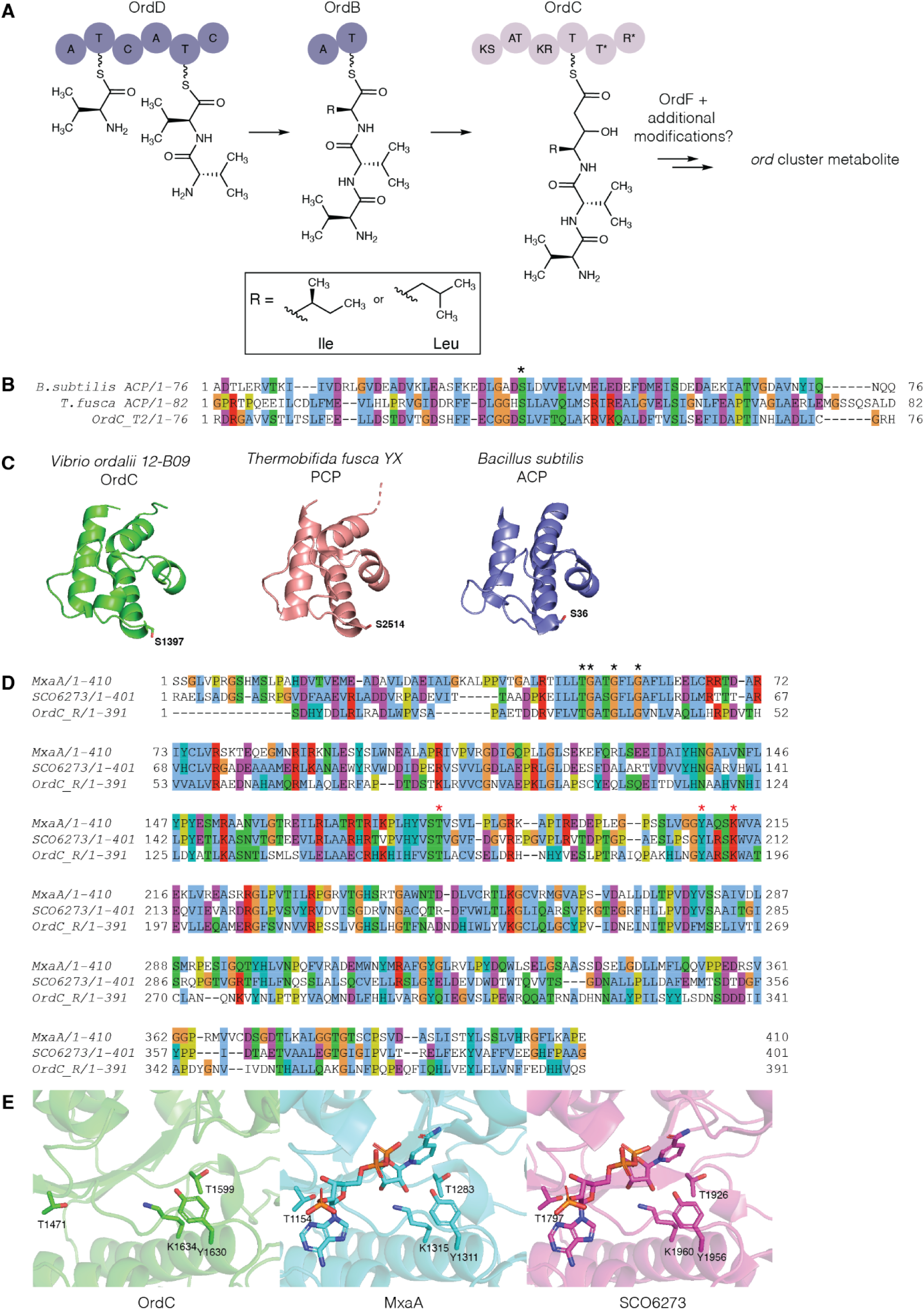
Bioinformatic analysis of the Ord proteins suggests biosynthetic roles. (**A**) Bioinformatic analysis informs domain substrate predictions *= predicted by antiSMASH to be incomplete. A = adenylation, T = thiolation, C = condensation, KS = ketosynthase, AT = acyltransferase, KR = ketoreductase, R = reductase. (**B**) Multiple sequence alignment of OrdC thiolation domain with carrier proteins from *Thermobifida fusca YX*(*77*) and *Bacillus subtilis*(*78*). Sequences were aligned in Geneious Prime v2023.2.1(*79*) using the MUSCLE algorithm with default parameters and colored in JalView(*80*) using the Clustal color scheme. Ppant-binding serine residues are indicated with an asterisk (*). (**C**) Comparison of an AlphaFold3(*81*) structure to crystal structures of carrier proteins from *T. fusca* (PDB ID: 8FX7)(*82*) and *B. subtilis* (PDB ID: 1HYB)(*83*) demonstrates conserved structure. (**D**) Multiple sequence alignment of OrdC with reductase domains of MxaA(*82*) and SCO6273. Sequences were aligned in Geneious Prime v2023.2.1(*79*) using the MUSCLE algorithm with default parameters and colored in JalView(*80*) using the Clustal color scheme. NAD(P)H binding residues are highlighted with a black asterisk (*) and catalytic residues are indicated with a red asterisk (*). (**E**) Comparison of an AlphaFold3(*81*) generated structure of OrdC to MxaA (PDB ID: 4U7W)(*82*) and SCO6273 (PDB ID: 8V1X)(*83*) crystal structures demonstrates conserved structure.

**Fig. S6.**
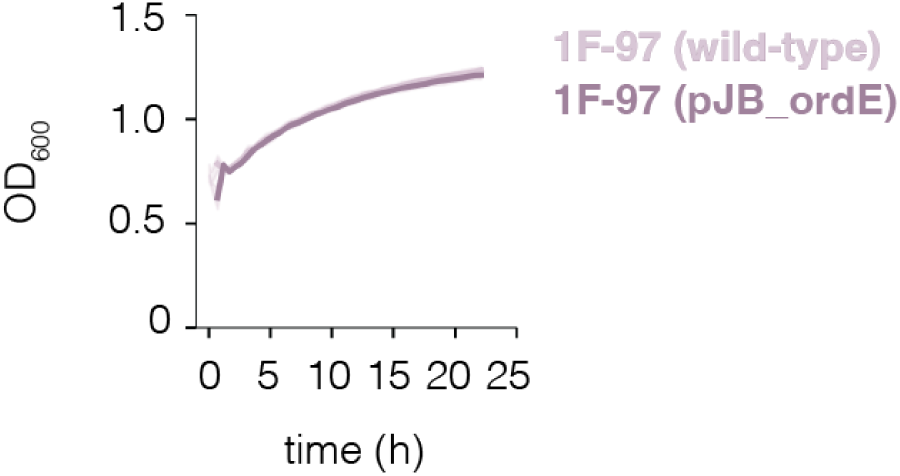
Presence of pJB_ordE does not incur growth defects in 1F-97. Growth measurements comparing 1F97 pJB_ordE to wild-type. Representative curve from three biological replicates, each with four technical replicates (mean with standard deviation).

**Fig. S7.**
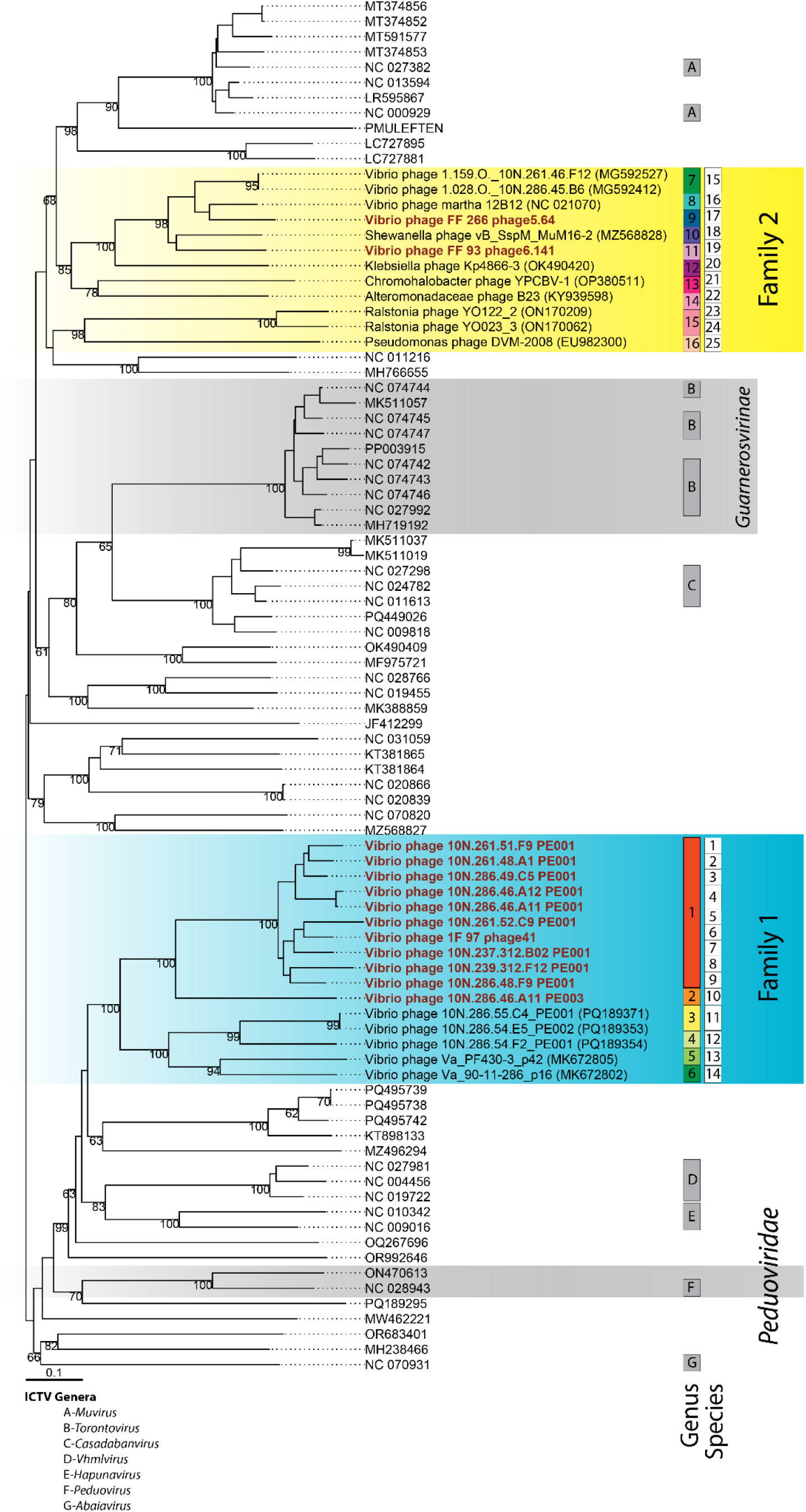
Amino acid-based whole-genome phylogeny determined with VICTOR for the induced phages and their relatives. Tree of the inducible prophages found either in the screen (phages 41 of *V. cyclitrophicus* 1F-97, 5.64 of *V. tasmaniensis* FF-266, 6.141 of *V. ordalii* FF-93) or in the subsequent search for phages similar to phage 41 (depicted in red) and blastp-determined(*54*) relatives in the *Caudoviricetes*. New phage families, as identified by VirClust(*65*), are marked, while genera and species—determined based on nucleotide identity using VIRIDIC(*66*)—are indicated with squares. Additionally, the official ICTV taxonomy (MSL40v.1) is shown in gray, encompassing the *Peduoviridae* family, the *Guanerosvirinae* subfamily, and seven genera. Pseudo-bootstrap values are displayed along the branches.

**Fig. S8.**
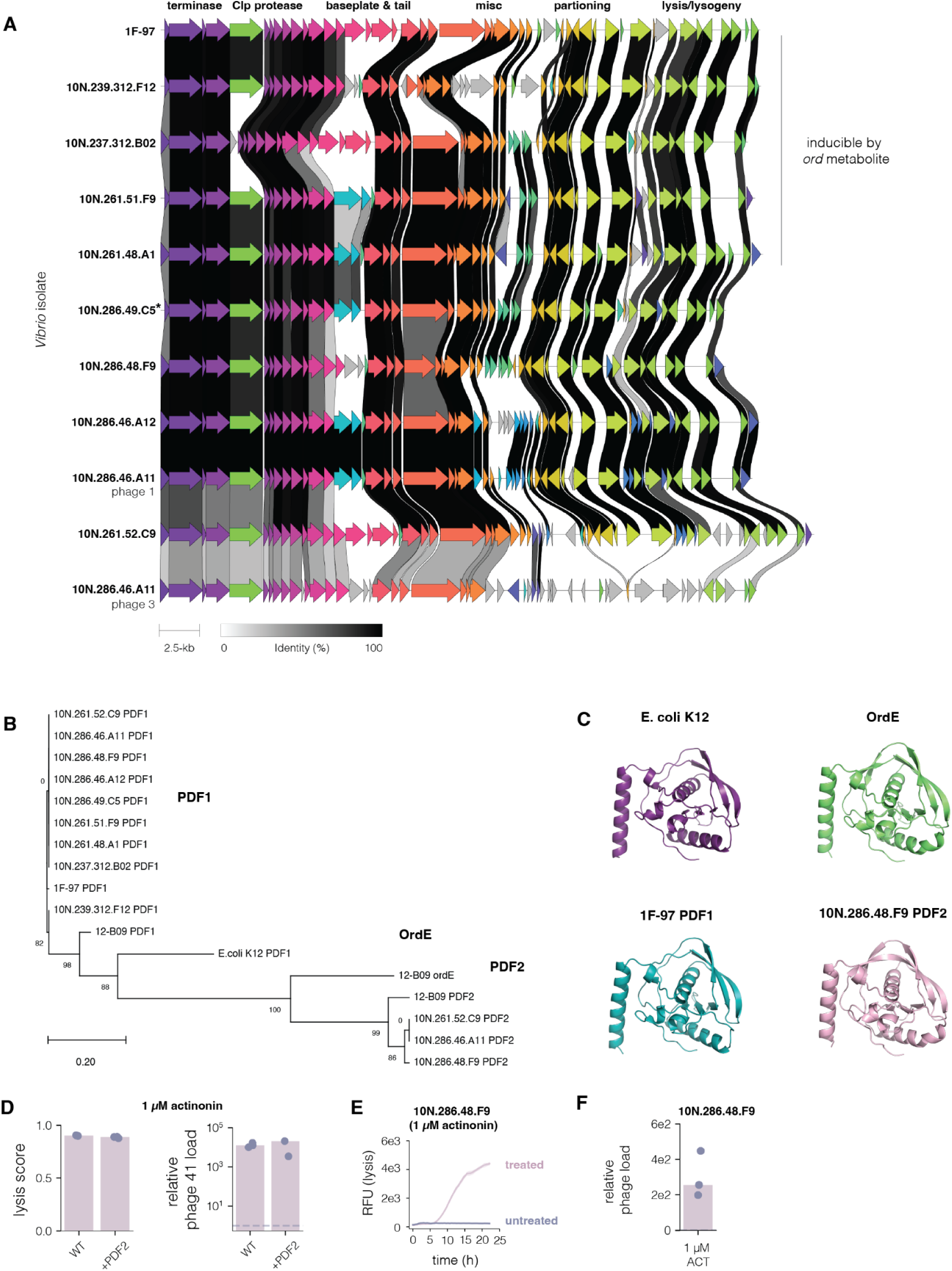
Alternative peptide deformylases enable specific resistance to the *ord* cluster metabolite. (**A**) Alignment (identity cutoff of 30%) of phage 41 and similar phages detected in other *Vibrio* hosts. (**B**) Protein tree for all peptide deformylases found in the *Vibrio* hosts from (A), 12-B09 (including OrdE), and the canonical peptide deformylase from *E. coli*. The tree was generated using MUSCLE (v5.3)(*57*) alignment of the PDF amino acid sequences, FastTree (v2.1.11)(*58*) with the WAG+CAT model, and visualized with MEGA X (v10.1)(*59*). (**C**) The protein structures were predicted using AlphaFold 3(*60*) for 12-B09’s OrdE, 1F-97 PDF1, and PDF2 from *Vibrio lentus* 10N.286.48.F9 to determine conservation of the structure compared to the predicted structure of PDF1 from *E. coli* K12. (**D**) Lysis scores and estimated degree of phage 41 induction upon actinonin treatment (1 µM) when 1F-97 expresses PDF2 from 10N.286.48.F9. (**E**) Lysis and (**F**) prophage induction of the prophage found in 10N.286.48.F9 that resembles phage 41 of 1F-97 upon actinonin treatment (1 µM). (D) Lysis score bars represent the mean of three biological replicates with individual replicates shown (mean of four technical replicates). (E) Representative curves from three biological replicates are shown from a 22-h time course performed in 384W microtitre plates, with four technical replicates per culture, represented as the mean with standard deviation. (D,F) qPCR bars represent the median of three biological replicates with individual replicates shown (each with three technical replicates, aggregated as the mean).

**Fig. S9.**
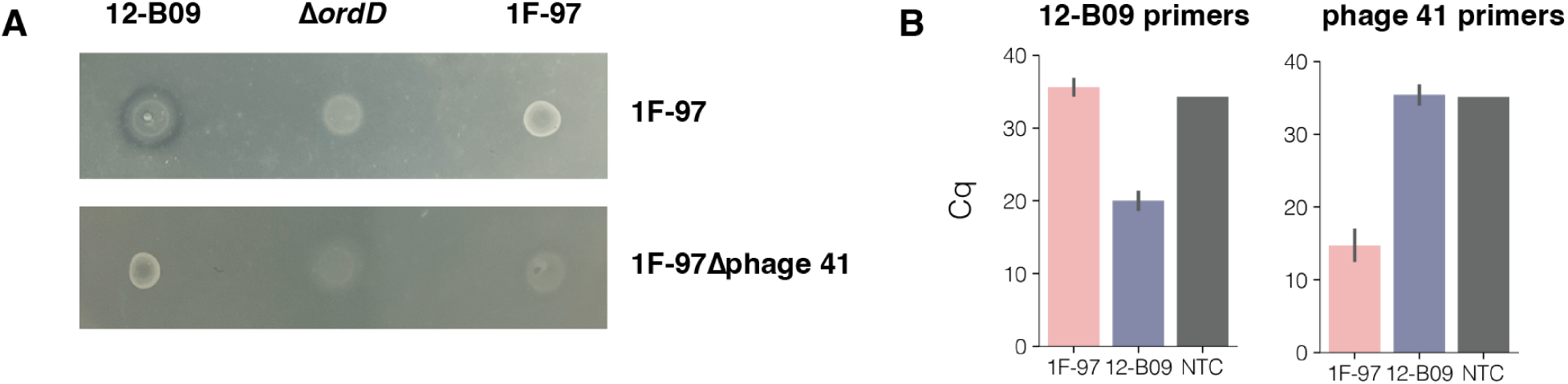
12-B09 requires NRPS *ordD* for successful interference competition. (**A**) Burkholder assay of 12-B09 (wild-type or Δ*ordD* mutant spots) against 1F-97 (wild-type or Δphage 41 top agar) with wild-type 1F-97 spots as a negative control. A dark ring around the spot demonstrates a zone of clearance where the target strain is inhibited. Representative drop spots are shown for each assay of three biological replicates, each with three technical replicates. (**B**) To determine whether 12-B09 was infected and lysogenized by phage 41 post-induction, the presence of phage 41 in single colonies (plated post co-culture in broth) were quantified using phage-specific primers. The bars represent mean C_q_ from 12-B09 colonies (n=44 in indigo) and 1F-97 colonies (n=5 in pink) shown with standard deviation and no template control (NTC, in grey) (n=1). The phage 41 primers demonstrated a 20.7 cycle-difference (fold-difference of 1.7 x 10^6^) where 12-B09 colonies resembled the NTC. Primers against the 12-B09 genome were included to confirm the strain identity of the colony.

**Fig. S10.**
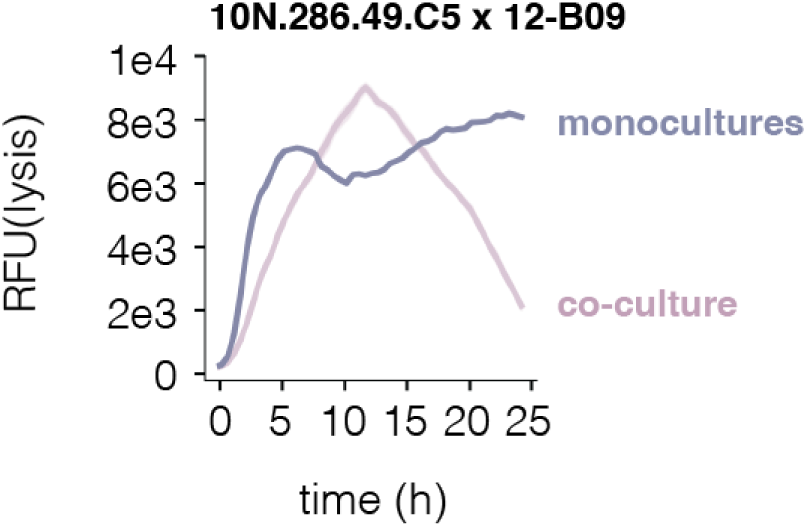
10N.286.49.C5 exhibits high baseline lysis in monoculture. Lysis curves to compare the sum of 12-B09 and 10N.286.49.C5 monocultures to their co-culture with (initial inocula of OD_600_ 0.05 for each). Corresponds to the hosts of phages resembling phage 41 in fig. 3E and fig. S8A. Representative lysis curves from three biological replicates are shown performed in 384W microtitre plates, with four technical replicates per culture, represented as the mean with standard deviation.

## Supplementary Tables

**Table S1.** Putative function of proteins encoded by the *ord* biosynthetic gene cluster in *Vibrio ordalii* 12-B09. Closest characterized homologs were identified through BLAST searches in the UniProtKB/Swiss-Prot database.

| Protein | Size (aa) | Annotation | Homolog origin | Proposed function | Accession ID | Identity (% aa) | Query coverage (%) |
| --- | --- | --- | --- | --- | --- | --- | --- |
| OrdA | 249 | 4'-phosphopantetheinyl (Ppant) transferase | *Salmonella enterica* subsp. *Enterica* serovar *Typhi* | Phosphopantetheinyl transferase | Q8Z8L9.1 | 32.6 | 73 |
| OrdB | 559 | NRPS | *Brevibacillus parabrevis* | NRPS | O30409.1 | 31.7 | 98 |
| OrdC | 1833 | PKS | *Mycobacterium tuberculosis* CDC1551 | PKS | P9WQE0.1 | 34.4 | 43 |
| OrdD | 2052 | NRPS | *Brevibacillus parabrevis* | NRPS | Q70LM5.1 | 99 | 29.7 |
| OrdE | 171 | Peptide deformylase | *Vibrio parahaemolyticus* RIMD 2210633 | Peptide deformylase | Q87I22.1 | 97 | 60.8 |
| OrdF | 275 | methyltransferase | *Corynebacterium urealyticum* DSM 7109 | methyltransferase | B1VEN4.1 | 36 | 38.2 |

**Table S2.**
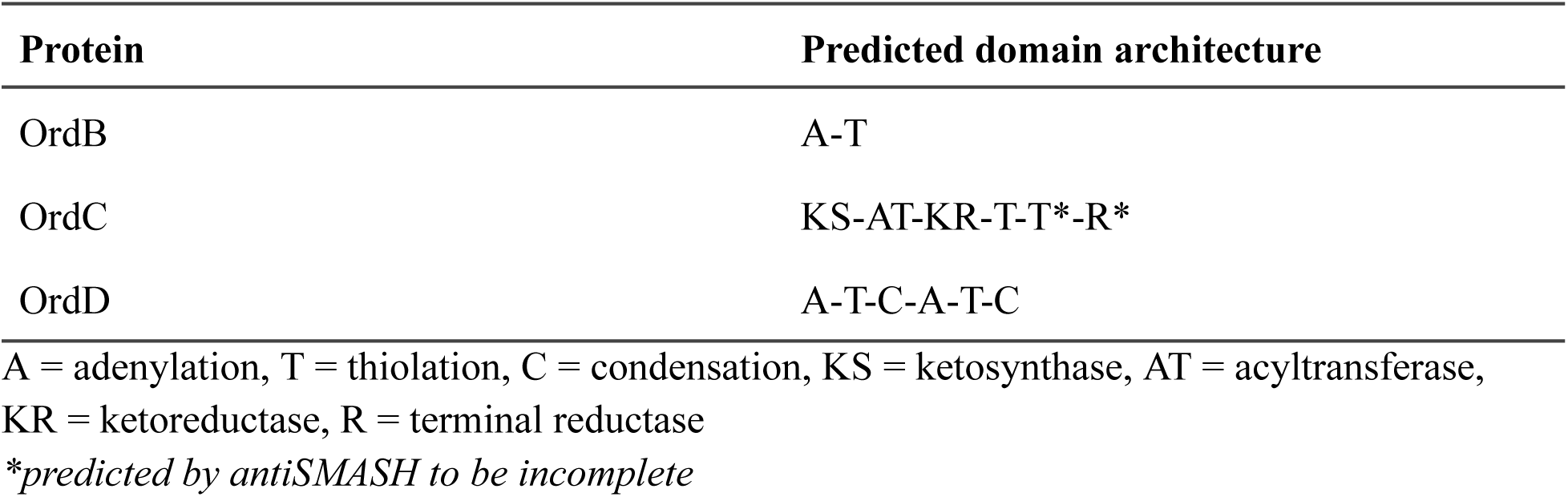
antiSMASH predicted domains of assembly line proteins OrdB, OrdC, and OrdD.

| **Protein** | **Predicted domain architecture** |
| --- | --- |
| OrdB | A-T |
| OrdC | KS-AT-KR-T-T*-R* |
| OrdD | A-T-C-A-T-C |

A = adenylation, T = thiolation, C = condensation, KS = ketosynthase, AT = acyltransferase, KR = ketoreductase, R = terminal reductase
*\*predicted by antiSMASH to be incomplete*

**Table S3.** Substrate predictions for the assembly line proteins OrdB, OrdC, and OrdD.

| **Protein** | **Specificity-conferring residues** | **antiSMASH (aS)** | **PRISM** | **Univ. of Maryland NRPS Predictor (UM)** |
| --- | --- | --- | --- | --- |
| OrdB | D-A-F-F-Y-G-C-V-W-K (aS)<br>D-A-F-A-Y-G-C-V (UM) | Leucine | Valine | Leucine, isoleucine, or valine |
| OrdC | Not applicable | Malonyl-CoA | Malonyl-CoA | No data |
| OrdD (1) | D-A-L-W-M-G-G-T-F-K (aS)<br>D-A-L-W-M-G-G-T (UM) | Valine | Phenylalanine | Valine |
| OrdD (2) | D-A-L-W-L-G-G-T-F-K (aS)<br>D-A-L-L-L-G-G-T (UM) | Valine | Valine | No data |

